# SOS1 and KSR1 modulate MEK inhibitor responsiveness to target resistant cell populations based on PI3K and KRAS mutation status

**DOI:** 10.1101/2022.12.06.519395

**Authors:** Brianna R. Daley, Heidi M. Vieira, Chaitra Rao, Jacob M. Hughes, Zaria M. Beckley, Dianna H. Huisman, Deepan Chatterjee, Nancy E. Sealover, Katherine Cox, James W. Askew, Robert A. Svoboda, Kurt W. Fisher, Robert E. Lewis, Robert L. Kortum

## Abstract

KRAS is the most commonly mutated oncogene. Targeted therapies have been developed against mediators of key downstream signaling pathways, predominantly components of the RAF/MEK/ERK kinase cascade. Unfortunately, single-agent efficacy of these agents is limited both by intrinsic and acquired resistance. Survival of drug-tolerant persister cells (DTPs) within the heterogeneous tumor population and/or acquired mutations that reactivate receptor tyrosine kinase (RTK)/RAS signaling can lead to outgrowth of tumor initiating cells (TICs) and drive therapeutic resistance. Here, we show that targeting the key RTK/RAS pathway signaling intermediates SOS1 or KSR1 both enhances the efficacy of, and prevents resistance to, the MEK inhibitor trametinib in *KRAS*-mutated lung (LUAD) and colorectal (COAD) adenocarcinoma cell lines depending on the specific mutational landscape. The SOS1 inhibitor BI-3406 enhanced the efficacy of trametinib and prevented trametinib resistance by targeting spheroid initiating cells (SICs) in *KRAS^G12/G13^*-mutated LUAD and COAD cell lines that lacked *PIK3CA* co-mutations. Cell lines with *KRAS^Q61^* and/or *PIK3CA* mutations were insensitive to trametinib and BI-3406 combination therapy. In contrast, deletion of the RAF/MEK/ERK scaffold protein *KSR1* prevented drug-induced SIC upregulation and restored trametinib sensitivity across all tested *KRAS* mutant cell lines in both *PIK3CA*- mutated and *PIK3CA* wildtype cancers. Our findings demonstrate that vertical inhibition of RTK/RAS signaling is an effective strategy to prevent therapeutic resistance in *KRAS*- mutated cancers, but therapeutic efficacy is dependent on both the specific KRAS mutant and underlying co-mutations. Thus, selection of optimal therapeutic combinations in *KRAS*-mutated cancers will require a detailed understanding of functional dependencies imposed by allele-specific KRAS mutations.

**Significance Statement:** We provide an experimental framework for evaluating both adaptive and acquired resistance to RAS pathway-targeted therapies and demonstrate how targeting specific RAS pathway signaling intermediates SOS1 or KSR1 enhanced effectiveness of and prevented resistance to MEK inhibitors in *KRAS*-mutated cancer cells with genotypic precision. The contribution of either effector was dependent upon the mutational landscape: SOS1 inhibition synergized with trametinib in *KRAS^G12/G13^*-mutated cells expressing WT PI3K but not in *KRAS^Q61^*-mutated cells or if *PIK3CA* is mutated. *KSR1* deletion inhibited MEK/ERK complex stability and was effective in cells that are unresponsive to SOS1 inhibition. These data demonstrate how a detailed understanding of functional dependencies imposed both by allele specific *KRAS* mutations and specific co-mutations facilitates the optimization of therapeutic combinations.

## Introduction

RAS proteins are encoded by three paralogs, KRAS, NRAS, and HRAS, which are, collectively, the most frequently mutated oncogene in cancer (1, 2). Among those paralogs, KRAS is the most commonly mutated, found predominantly in pancreas adenocarcinoma (PDAC) (95%), lung adenocarcinoma (LUAD) (30-40%), and colorectal adenocarcinoma (COAD) (45-50%) (3). KRAS is commonly mutated at one of three mutational hotspots, G12, G13, or Q61 (4); mutation of one of these sites alters KRAS GTP/GDP cycling leading to increased KRAS-GTP loading and hyperactivation of downstream effectors including the pro-proliferative RAF/MEK/ERK kinase cascade. The RAF/MEK/ERK kinase cascade is the critical driver of proliferation in *KRAS*-mutated cancers (5–9), and multiple small molecule inhibitors of each kinase have been evaluated in *KRAS*-mutated cancers (10). Of these, the MEK inhibitors trametinib and selumetinib are among the most promising agents (11, 12). Unfortunately, single-agent treatment with MEK inhibitors is largely ineffective in *KRAS*-mutated cancers due to both intrinsic (adaptive) and acquired resistance. Intrinsic resistance occurs due to the presence of pre-existing mechanisms that render tumor cells insensitive to that specific therapeutic intervention (13). For MEK inhibitors, intrinsic resistance is driven both by relief of ERK-dependent negative feedback of RTK−SOS−WT RAS−PI3K signaling (14–18) and compensatory ERK reactivation (5, 19, 20). Thus, either broad inhibition of RTK rebound signaling and/or deep inhibition of MEK/ERK signaling may be required to enhance the efficacy of MEK inhibitors to treat *KRAS*-mutated cancers (18, 21, 22).

Even if one is able to overcome intrinsic/adaptive resistance, treatment failure can also occur via acquired resistance, where resistance-conferring mutations, phenotypes, or shifts in oncogenic signaling that occur under selective pressure lead to tumor outgrowth after an initial period of drug responsiveness (13, 21). *KRAS*-mutated cancer cells treated with MEK inhibitors are capable of surviving targeted treatments by entering a near quiescent state (5, 23), becoming drug-tolerant persister (DTP) cells (21). DTPs exhibit subpopulations of highly plastic cells with altered metabolic and drug efflux properties (21, 24) also known as tumor initiating cells (TICs) *in vivo* or spheroid initiating cells (SICs) *in vitro*. TICs/SICs exhibit stem-like properties, can self-renew and divide asymmetrically to give rise to additional cell types within the tumor, and may represent the sanctuary population within the bulk tumor responsible for treatment failure and recurrence (25, 26). In colorectal cancer, MEK inhibition may increase the TIC population through promotion of stem-like signaling pathways (27) and targeting TIC emergence may be required to circumvent acquired resistance.

*KRAS*-mutated cancers are addicted to RTK/RAS signaling, and combination therapeutic strategies that vertically inhibit RTK/RAS/effector signaling represents an attractive approach to limiting MEK inhibitor induced rebound RTK−PI3K signaling and compensatory ERK reactivation in *KRAS*-mutated cancers (5, 14–20). Upstream of RAS, the RAS guanine nucleotide exchange factors (RasGEFs) SOS1/2 regulate RTK-stimulated RAS activation and represent a key ‘stoichiometric bottleneck’ for RTK/RAS pathway signaling (28). We previously showed that *SOS2* deletion synergized with trametinib to inhibit anchorage-independent survival in *KRAS*-mutated cancer cells (18), but only in cells with WT *PIK3CA*. While no SOS2 inhibitors have been developed to date, multiple groups have developed SOS1 inhibitors with the goal of using these to treat RTK/RAS mutated cancers (29–35). The most well characterized SOS1 inhibitor, BI-3406, has modest single-agent efficacy in *KRAS*-mutated cells but enhanced the efficacy of the MEK inhibitor trametinib in *KRAS*-mutated xenografts (32). BI-3406 activity is RAS codon-specific, killing cells harboring *KRAS^G12^* and *KRAS^G13^* mutations that are dependent upon activation by GEFs, but not cells harboring *KRAS^Q6^*^1^ mutations. Mutation of Q61 dramatically reduces intrinsic hydrolysis compared to either G12 or G13 mutations, promoting GEF-independent signaling (36, 37).

Downstream of RAS, Kinase Suppressor of RAS 1 (KSR1) is a molecular scaffold for the RAF/MEK/ERK kinase cascade that controls the intensity and duration of ERK signaling to dictate cell fate (38–40). While KSR1 is required for mutant RAS-driven transformation (38) and tumorigenesis (41), it is dispensable for normal growth and development (41–43).

Here, we demonstrate that enhanced efficacy of, and delayed resistance to, the MEK inhibitor trametinib can be achieved through vertical inhibition of RTK/RAS signaling in *KRAS*-mutated cancer cells. However, we further found that the optimal co-targeting strategy is dependent both on the specific KRAS allelic mutation and the presence of *PIK3CA* co-mutations. In *KRAS^G12^*- and *KRAS^G13^*-mutated LUAD and COAD cells, the SOS1 inhibitor BI-3406 synergistically enhanced trametinib efficacy and prevented the development of trametinib resistance by targeting SICs. These effects were lost in *KRAS^Q6^*^1^-mutated cells or if *PIK3CA* is mutated. In contrast, *KSR1* knockout (KO) limited TIC/SIC survival and trametinib resistance in both *KRAS^Q61^*-mutated cells and in *KRAS*-mutated COAD cells with *PIK3CA* co-mutations in an ERK-dependent manner. Thus, selection of optimal therapeutic combinations in *KRAS*-mutated cancers will require a detailed understanding of functional dependencies imposed by allele-specific KRAS mutations.

## Results

### SOS1 inhibition synergizes with trametinib to prevent rebound signaling in *KRAS^G12^/PIK3CA^WT^*-mutated LUAD cells

BI-3406 is a potent, selective SOS1 inhibitor previously shown to reduce 3D proliferation of *KRAS^G12/G13^*-mutated, but not *KRAS^Q61^*- mutated, cell lines as a single agent and to enhance the efficacy of trametinib in *KRAS*- mutated xenografts (32). To characterize the extent to which BI-3406 enhances the effectiveness of trametinib, we treated a panel of 3D spheroid cultured *KRAS*-mutated LUAD cell lines with increasing doses of BI-3406 and/or trametinib in a 9x9 matrix of drug combinations and assessed for synergistic killing after 96 hours by Bliss Independence (Fig. 1A). We found that in *KRAS^G12^*-mutated cell lines H727 (G12V), A549 (G12S), and H358 (G12C), SOS1 inhibition markedly enhanced the efficacy of trametinib at or below the EC_50_ for trametinib as assessed by significant reductions in the EC_50_ of trametinib (Fig. 1 A and *SI Appendix, Fig. S1*) and showed a high excess over Bliss across the treatment matrix indicative of drug-drug synergy (Fig. 1B).

**Fig 1.**
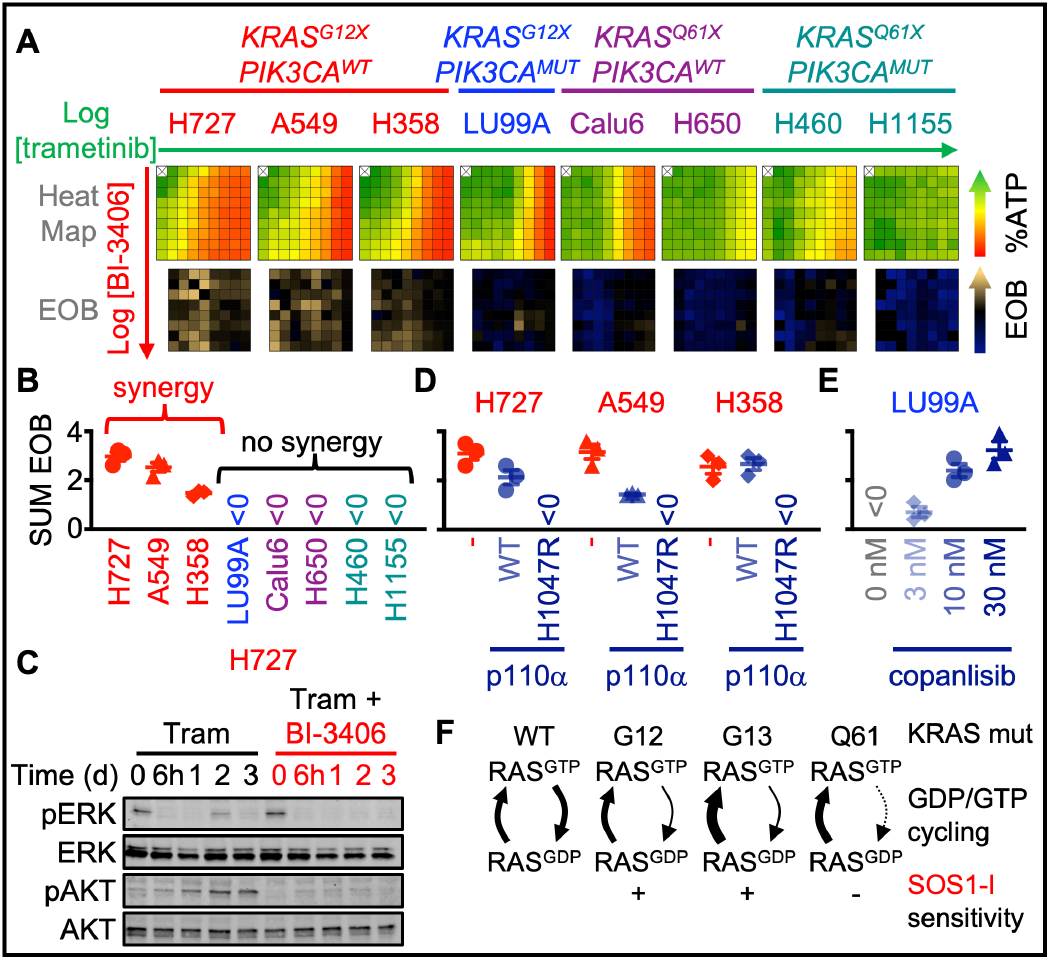
MEK and SOS1 inhibition synergize to prevent rebound signaling in *KRAS^G12^*/*PIK3CA^WT^*-mutated LUAD cells. *(A)* Heat map of cell viability (top) and excess over Bliss (EOB, bottom) for the indicated KRAS-mutated LUAD cell lines treated with increasing (semilog) doses of trametinib (10^-10.5^ – 10^-7^), BI-3406 (10^-9^ – 10^-5.5^) or the combination of trametinib + BI-3406 under 3D spheroid culture conditions. The KRAS and PIK3CA mutational status of each cell line is indicated. Data are the mean from three independent experiments, each experiment had three technical replicates. *(B-D)* The sum of excess over Bliss for the 9×9 matrix of cells treated with trametinib + BI3406 from *A (B)*, *KRAS^G12^*/*PIK3CA^WT^* cells expressing a WT or H1047R mutant p110α catalytic subunit *(C)*, or KRAS^G12^/PIK3CA^mut^-mutated LUAD cells treated with increasing doses of copanlisib *(D)*. EOB > 0 indicates increasing synergy. *(E)* Western blots of WCLs of 3D spheroid cultured H727 cells treated with trametinib (10 nM) ± BI-3406 (300 nM) for the indicated times. Western blots are for pERK, ERK, pAKT (Ser 473), AKT. *(F)* GDP/GTP RAS cycling in different KRAS mutations with the proposed SOS1 inhibitor sensitivities.

As a single agent, the effectiveness of trametinib is blunted by rapid induction of RTK/PI3K signaling followed by rebound ERK activation due, in part, to loss of ERK-dependent negative feedback signaling (14, 22, 44). In *KRAS^G12^*-mutated H727 and A549 cells, BI-3406 reduced the trametinib-induced increase in PI3K/AKT activation in a time (Fig. 1 C, and *SI Appendix, Fig. S2 A*) and dose (*SI Appendix, Fig. S2 B*)- dependent manner. BI-3406 further enhanced trametinib-induced inhibition of pERK/pRSK and limited rebound ERK activation (Fig. 1 C, and *SI Appendix, Fig. S2 A and B*). These data suggest that SOS1 inhibition blocked PI3K-dependent adaptive resistance to MEK inhibitors and decreased the effective dose at which trametinib blocked ERK signaling in *KRAS^G12^*-mutated LUAD cells.

Consistent with the hypothesis that RTK/PI3K signaling drives adaptive resistance to trametinib, SOS1 inhibition did not synergize with trametinib (Fig. 1 A and B and *SI Appendix, Fig. S1*) or inhibit rebound signaling (*SI Appendix, Fig. S2*) in *KRAS^G12^*-mutated LU99A cells that harbor an activating *PIK3CA* co-mutation. To determine the extent to which mutational activation of PI3K/ATK signaling was sufficient to limit the ability of SOS1 inhibition to enhance the efficacy of trametinib, we expressed either a wild-type or an activated form of the p110α catalytic subunit (p110α^H1047R^) in *KRAS^G12^*-mutated cell lines H727 (G12V), A549 (G12S), and H358 (G12C) lacking *PIK3CA* mutations that were previously shown to be sensitive to combined trametinib/BI- 3406 treatment (*SI Appendix, Fig. S3 A*). Expression of p110α^H1047R^, but not wild type p110α, abrogated synergy between trametinib and BI-3406 in all three cell lines (Fig. 1 C and *SI Appendix, Fig. S3 A and B*). To determine the extent to which activated PI3K signaling was necessary to limit trametinib/BI-3406 synergy in LU99A cells, we treated LU99A cells with increasing doses of the PI3K inhibitor copanlisib in combination with increasing doses of BI-3406 and/or trametinib in a 9x9 matrix of drug combinations. We found that PI3K inhibition caused a dose-dependent sensitization of the *KRAS^G12C^*/*PIK3CA* mutated LU99A cells to the combination of trametinib and BI-3406 (Fig. 1 D and *SI Appendix, Fig. S4*). These data suggest a role for SOS1 in adaptive resistance to trametinib in a PI3K-dependent manner.

SOS1 inhibition also failed to synergize with trametinib (Fig. 1 A and B and *SI Appendix, Fig. S1*) or alter activation of RAF/MEK/ERK or PI3K/AKT signaling in a time- or dose-dependent manner (*SI Appendix, Fig. S4*) in *KRAS^Q61^*-mutated LUAD cell lines regardless of *PIK3CA* mutation status, confirming previous studies where SOS1 inhibition is only effective in *KRAS*-mutated cancer cells where KRAS cycles between the GTP and GDP-bound state (Fig. 1 F) and (32). To confirm the effects of SOS1 inhibition were due to inhibition of RAS signaling, we further assessed the relative levels of active RAS in NT and *SOS1* KO cells treated with BI-3406 for 24 hours (*SI Appendix, Fig. S5*). Indeed, *SOS1* KO or BI-3406 treatment decreased the levels of active RAS only in *KRAS^G12^*-mutated cells where mutant KRAS actively cycles between an active and inactive state, but not in *KRAS^Q61^*-mutated cells where mutant KRAS cycles independently of RAS-GEF activity (Fig. 1 F and *SI Appendix, Fig. S5*). BI-3406 treatment had no additional effect on the levels of active RAS in *SOS1* KO cells, confirming the specificity of BI-3406 toward SOS1 (*SI Appendix, Fig. S5*).

### Combination therapy with MEK and SOS1 inhibition targets trametinib-induced SIC outgrowth

Single agent therapy with EGFR Tyrosine Kinase Inhibitors increases SIC populations in NSCLC (45). We found that MEK inhibitors similarly expand SIC populations in *KRAS*-mutated LUAD cells (Fig. 2). Aldehyde dehydrogenases (ALDH) are enzymes that oxidize aldehydes (46, 47) and have been proposed as a functional marker of tumor initiating cells that detoxify the effects of therapy-induced oxidative stress to promote survival of LUAD TICs (48–51). Most NSCLC cell lines have a sub- population of cells exhibiting elevated ALDH activity, although the absolute abundance of ALDH^+^ cells between different NSCLC cell lines does not directly correlate with differences in clonogenicity between cell lines of distinct origins (50). However, within a given LUAD cell line isolated ALDH^+^ cells show increased clonogenicity (50, 52–54) and resistance to conventional and targeted therapies (54, 55), and conversely depletion or inhibition of ALDH reduces clonogenicity (47, 48, 54). Further, the frequency of cells showing increased ALDH activity increases in drug tolerant persister cells that survive EGFR-targeted therapies in *EGFR*-mutated LUAD (47, 55–57). We thus first assessed the extent to which MEK inhibition changed the frequency of ALDH^+^ cells as a measure of cells responding to increased oxidative stress in *KRAS^G12^*-mutated *PIK3CA^WT^* cells. MEK inhibitors trametinib and selumetinib caused a >3-fold increase in the frequency of ALDH^+^ cells in H727, A549, and H358 cells (Fig. 2 A and *SI Appendix, Fig. S6 A*). We used an Extreme Limiting Dilution Analysis (ELDA) in H727, A549, and H358 cells to assess spheroid growth in 96-well ultra-low attachment plates and determine the frequency of SICs. ELDAs were performed 72 hours after MEK inhibition with trametinib or selumetinib and assessed after 7-10 days for SIC outgrowth. ELDA results demonstrated a 2-3-fold significant increase in SIC frequency in MEK-inhibitor treated cells in comparison to untreated cells (Fig. 2 B).

**Fig. 2.**
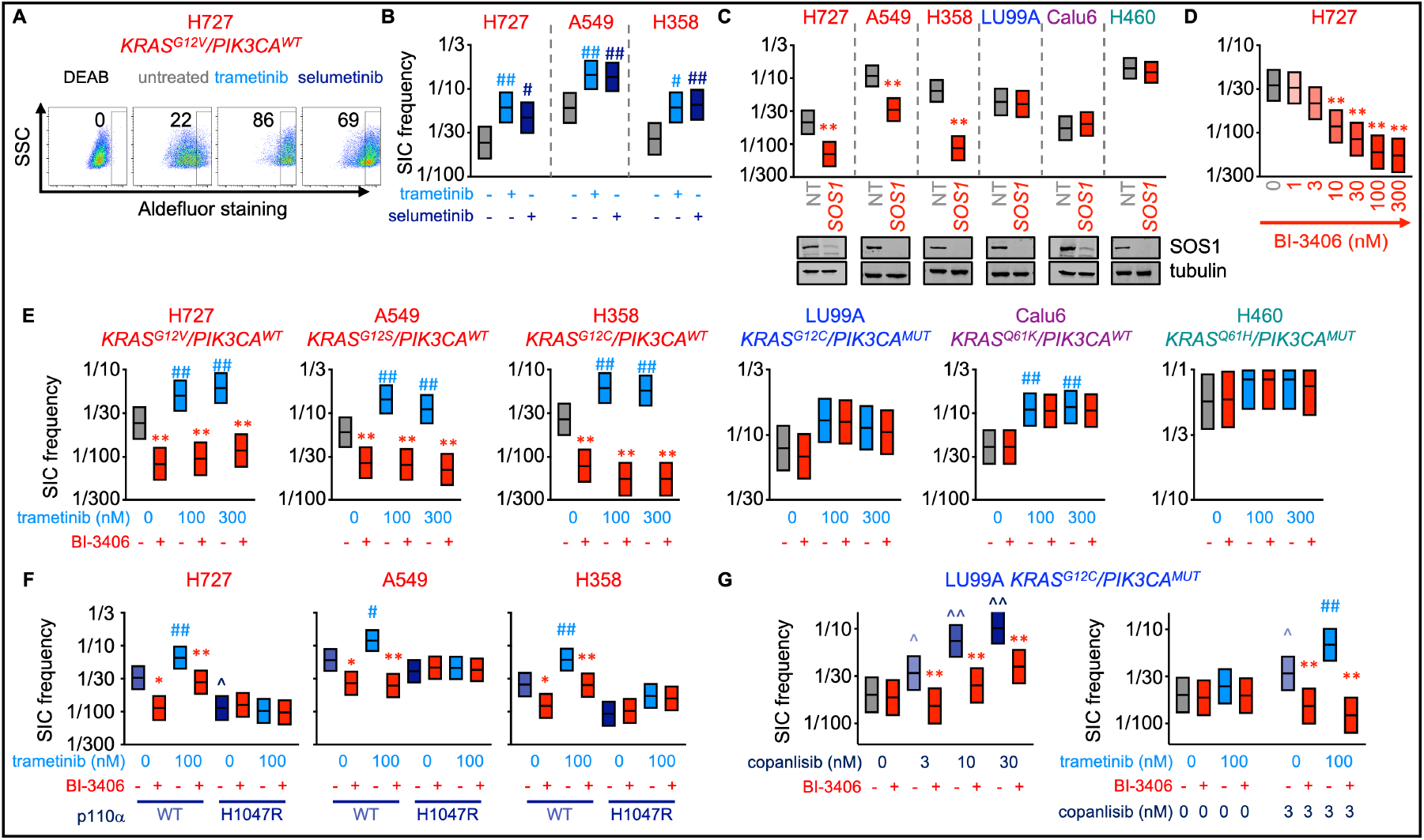
SOS1 inhibition prevents trametinib-induced SIC outgrowth. *(A)* Aldefluor staining for ALDH enzyme activity in DEAB negative control (DEAB), untreated H727 cells, or H727 cells treated with 100 nM trametinib or selumetinib for 72 hours. *(B-G)* SIC frequency from *in situ* ELDAs of the indicated cell lines pre-treated with 100 nM trametinib or selumetinib for 72 h *(B)*, cells where *SOS1* has been knocked out vs. non-targeting controls (C), H727 cells treated with the indicated doses of BI-3406 (D), cells pre-treated with trametinib for 72 h to upregulate TICs and then left untreated or treated with BI-3406 *(E)*, cells expressing WT or H1047R mutant p110α pre-treated with trametinib for 72 hours to upregulate TICs and then left untreated or treated with BI-3406 *(F)*, LU99A cells treated with the indicated dose of copanlisib alone (left) or pre-treated with 100 nM trametinib ± the indicated dose of copanlisib for 72 h to upregulate TICs and then left untreated or treated with BI-3406 *(G)*. # p<0.05 vs untreated; ## p<0.01 vs. untreated for TIC upregulation by MEK inhibitor treatment vs. untreated controls. * p < 0.05 vs untreated; ** p<0.01 for TIC inhibition by BI-3406 treatment compared to untreated controls; ^ p<0.05, ^^ p<0.01 vs untreated for copanlisib treated cells. Data are representative of three independent experiments.

SOS1 inhibition was effective in blocking adaptive resistance and enhancing the efficacy of trametinib (Fig. 1), leading us to assess whether *SOS1* KO would be able to kill persister cells in *KRAS*-mutated LUAD cells. Compared to NT control cells, *SOS1* KO caused a 3-5-fold significant decrease in SIC frequency in *KRAS^G12^*-mutated/*PIK3CA^WT^*cells (Fig. 2 C). SOS1 inhibition with BI-3406 decreased SIC frequency in a dose-dependent manner, with the greatest effect found at 300 nM, in H727 and A549 cells (Fig. 2 C and *SI Appendix, Fig. S6 B*). Since SOS1 was required for SIC survival, we hypothesized that SOS1 inhibition would also limit the survival of the increased SICs present following MEK inhibition. To test this hypothesis, we pre-treated cells with two doses of trametinib (Fig. 2E) or selumetinib (*SI Appendix, Fig. S6 C*) and used these cells to determine the extent to which SOS1 inhibition could limit survival of MEK inhibitor-induced SICs. We found that in H727, A549, and H358 cells, SOS1 inhibition targeted and significantly decreased the MEK-induced increase in SIC frequency, causing a 5-10-fold significant decrease in SICs in MEK-inhibitor treated cells (Fig. 2E and *SI Appendix, Fig. S6 C*). To determine the extent to which SOS1 is necessary for the MEK-inhibitor induced increase in SIC frequency, we pre-treated NT control or *SOS1* KO cells with trametinib for 72 hours and assessed SIC frequency by *in vitro* ELDA. *SOS1* KO both decreased the intrinsic frequency of SICs and inhibited trametinib-induced SIC outgrowth (*SI Appendix, Fig. S7*), further supporting that SOS1 inhibition directly targeted SIC outgrowth. Further, SOS1 inhibition showed no added benefit in *SOS1* KO cells (*SI Appendix, Fig. S7 B*), confirming the specificity of SOS1 inhibition in limiting SIC survival. These findings support our hypothesis that BI-3406 can be used to enhance the efficacy of trametinib and prevent the development of resistance in the presence of *KRAS^G12/G13^*-mutated LUAD cells without a *PIK3CA* mutation.

*SOS1* KO and drug sensitivity is dependent upon the mutational profile of LUAD cells. *SOS1* KO had no effect on SIC frequency in *KRAS^G12^/PIK3CA^mut^* (LU99A) cells or *KRAS^Q61^*-mutated cells that are either *PIK3CA* wildtype (Calu6) or *PIK3CA* mutant (H460) (Fig. 2C). In *KRAS^Q61K^*/*PIK3CA^WT^* Calu6 cells, trametinib increased SIC frequency 2-3-fold, however, trametinib did not cause a significant increase in SICs in LUAD cells harboring a *PIK3CA* mutation (LU99A, H460) (Fig. 2E). These data suggest that trametinib-induced RTK-PI3K signaling, regulated by SOS1, may drive SIC outgrowth.

Similar to the regulation of trametinib/BI-3406 synergy by PI3K pathway mutation status observed in Fig. 1, mutational activation of PI3K signaling was both necessary and sufficient to limit both trametinib-induced changes in SIC frequency and to limit SOS1-dependent regulation of SICs. Trametinib pre-treatment did not increase the frequency of TICs in H727, A549, and H358 (*PIK3CA*^WT^) cells expressing an activated form of the p110α catalytic subunit (p110α^H1047R^), and the SICs in these cells were insensitive to SOS1 inhibition (Fig. 2 F and *SI Appendix, Fig. S8*). In LU99A cells harboring a *PIK3CA* mutation, the PI3K inhibitor copanlisib caused a dose-dependent increase in SIC frequency and restored the sensitivity of cells to BI-3406 (Fig. 2G). Further, copanlisib/trametinib co-treatment increased the SIC frequency in LU99A cells, which was blocked by the SOS1 inhibitor BI-3406 (Fig. 2G). These data suggest that SOS1 regulates SICs in a PI3K-dependent manner.

### *KSR1* KO restores trametinib responsiveness and inhibits SIC survival in *KRAS/PIK3CA* co-mutated LUAD cells

The RAF/MEK/ERK scaffold protein, KSR1, is a positive regulator of ERK-dependent signaling in *RAS*-mutant cancers, but dispensable to the growth of untransformed cells and could therefore be a promising therapeutic target downstream of oncogenic RAS (38, 40, 58). Structural analysis reveals that trametinib binds to the MEK-KSR complex (59). In *KRAS^Q61^*/*PIK3CA^mut^* cells, RAS cycles independently of SOS and SOS1 inhibition does not synergize with trametinib (Fig. 1A and B) or suppress SICs (Fig. 2E). We sought to determine whether inhibition of signaling downstream of RAS via KSR1 disruption affects SIC survival and trametinib sensitivity in *KRAS^Q61H^*/*PIK3CA^mut^*H460 LUAD cells. CRISPR/Cas9-mediated knockout of *KSR1* reduced TIC-frequency 4-fold by *in vivo* ELDA, demonstrating that KSR1 regulates the TIC populations in H460 cells (Fig. 3A). *KSR1* KO sensitized H460 cells to trametinib and selumetinib in a dose-dependent manner under both 2D (adherent) and 3D (spheroid) culture conditions (Fig. 3B and C and *SI Appendix, Fig. S9*). To directly determine whether *KSR1* KO enhances trametinib-induced killing of H460 cells, we assessed loss of membrane integrity that is associated with cell death using a CellTox Green Cytotoxicity Assay. Trametinib caused a dose-dependent increase in CellTox green staining, which occurred at lower doses and a greater overall magnitude of staining on a per cell basis in *KSR1* KO cells compared to either NT controls or *KSR1* KO cells expressing a KSR1 transgene (Fig. 3C). In trametinib-treated cells, *KSR1* KO both inhibited compensatory ERK/RSK/S6 reactivation (Fig. 3D and *SI Appendix, Fig. S10 A*) and synergized to inhibit ERK/RSK/S6 signaling in a dose-dependent manner (Fig. 3E and *SI Appendix, Fig. S10B*).

**Fig. 3.**
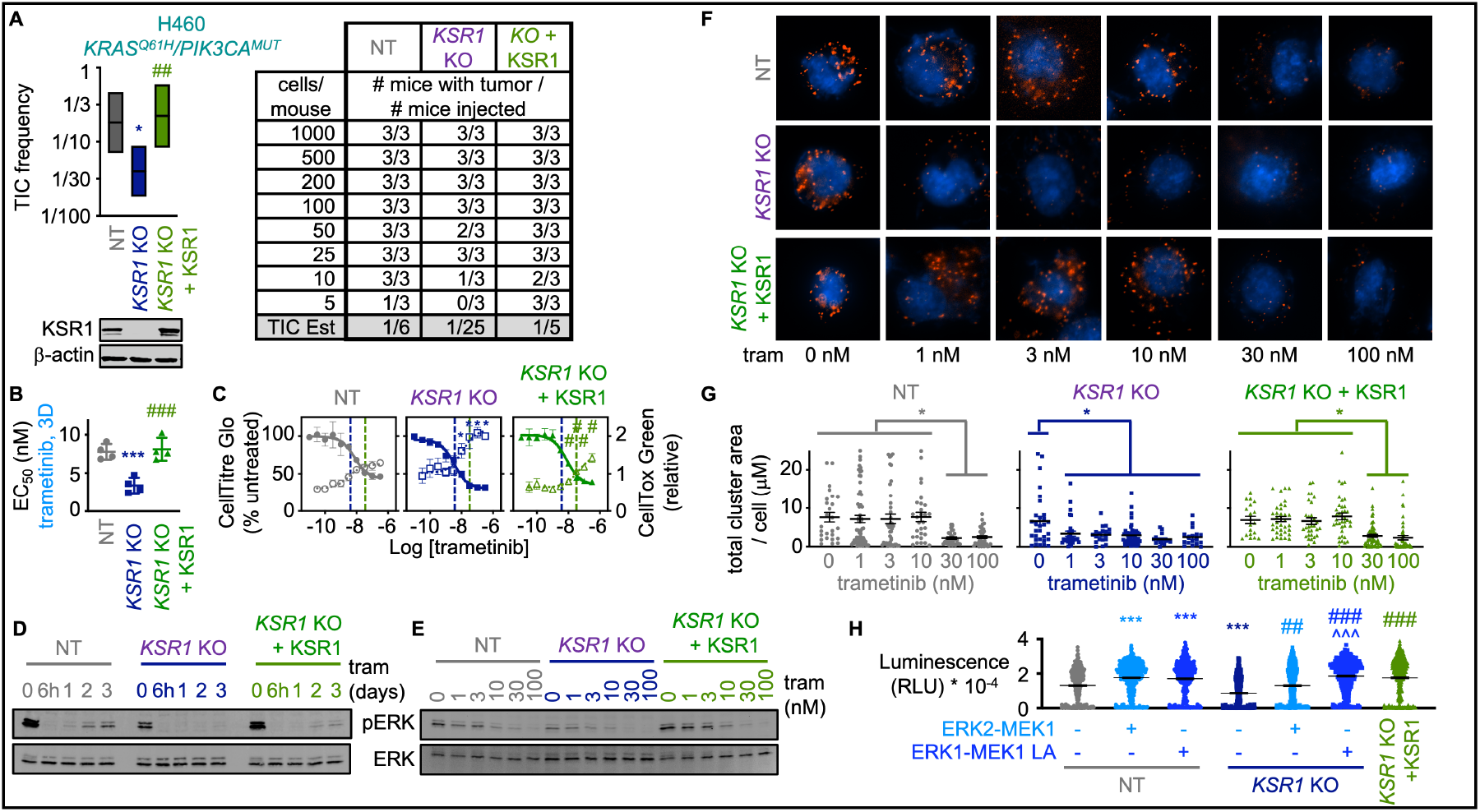
*KSR1* KO inhibits tumor initiating cell (TIC) survival and enhances sensitivity to trametinib in *KRAS^Q61^*-mutated LUAD cells. *(A) In vivo* limiting dilution analysis data showing TIC frequency in H460 (*KRAS^Q61H^/PIK3CA^MUT^*) cells. The indicated numbers of cells were injected into the shoulder and flank of NCG mice (Charles River). Tumors were scored at 30 days. *(B-C)* EC_50_ values *(B)* and trametinib dose-response curve indicating % cell viability (GellTitre Glo, left axis) and relative cell death (CellTox Green, right axis) *(C)* for H460 cells treated with the indicated concentrations of trametinib in anchorage-independent (3D) conditions for 72 hours. Vertical dashed lines show the intersection of viability and death curves for each population, *KSR1* KO or addback lines are shown for comparison. *(D-E)* Western blots of WCLs from H460 cells treated with trametinib (100 nM) for the indicated times *(D)* or with the indicated dose of trametinib for 24 hours *(E).* Western blots are for pERK and ERK. *(F)* Proximity ligation assay assessing MEK-ERK complex stability in H460 cells treated with the indicated dose of trametinib for 24 h; red= MEK-ERK complexes, blue = DAPI. *(G)* Quantification of the total area of MEK-ERK clusters from PLA assay in F. Each individual point represents a cell, data are quantified from >20 cells from three fields from three independent experiments. *(H)* Single cell colony forming assays for NT versus *KSR1* KO H460 cells expressing the indicated ERK2-MEK1 fusion proteins. Cells were single cell plated in non-adherent conditions, and colony formation was scored at 14 days by CellTitre Glo. Each individual point represents a colony. Western blotting for KSR1 and β-actin in each cell population are shown in A. *p<0.05; *** p<0.001 vs non-targeting controls; # p<0.05, ## p<0.01, ### p<0.001 vs. *KSR1* KO.

Since KSR1 is a scaffold protein for the RAF/MEK/ERK complex (38, 42), we used a MEK/ERK proximity ligation assay (20, 60, 61) to assess the extent to which *KSR1* KO enhanced trametinib responses by uncoupling MEK from ERK. In NT H460 cells, both the size and number of MEK/ERK complexes were significantly reduced at trametinib doses ≥ 30 nM leading to a marked decrease in the overall area of MEK/ERK clusters per cell (Fig. 3 F and G and *SI Appendix, Fig. S11*). While *KSR1* KO alone did not alter MEK/ERK interactions, *KSR1* KO led to a greater than 30-fold reduction in the dose of trametinib required to inhibit MEK/ERK complexes *in situ* (Fig. 3 F and G and *SI Appendix, Fig. S11*), which was rescued by ectopic KSR1 expression. To determine whether bypassing KSR1-dependent MEK-ERK scaffolding could restore clonogenicity in *KSR1* KO cells, we ectopically expressed either a WT ERK2-MEK1 fusion protein or a ERK2-MEK1 fusion with an inactivating mutation in the MEK nuclear export sequence (ERK2-MEK1 LA) that is transforming in fibroblasts (62). Single cell clonogenicity was used as an *in vitro* measure of SICs rather than *in situ* ELDA as H460 cells demonstrate a high (>50%) SIC frequency by *in situ* ELDA. Similar to the reduction in TICs observed by *KSR1* KO *in vivo* (Fig. 3A), *KSR1* KO significantly reduced single cell clonogenicity of H460 cells, which was rescued by ectopic KSR1 (Fig. 3H and *SI Appendix, Fig. S12*).

Ectopic expression of the ERK2-MEK1 fusion similarly restored H460 clonogenicity in *KSR1* KO cells, which was further increased by the transforming ERK2-MEK1 LA construct. These data demonstrate that inhibition of signaling distal to RAS, via *KSR1* KO, depletes SICs and enhances trametinib responsiveness in *KRAS^Q61H^/PIK3CA^MUT^*H460 cells via loss of KSR1 scaffolding of the MEK/ERK complex.

### In COAD, *KSR1* KO prevents trametinib-induced SIC increase regardless of PIK3CA mutational status

While *PIK3CA*/*KRAS* co-mutations are relatively rare in LUAD, they commonly co-occur in COAD, with approximately one third of *KRAS*- mutated COADs harboring co-existing *PIK3CA* mutations. We sought to test the extent to which the *KRAS/PIK3CA* genotype sensitivity to SOS1 and *KSR1* ablation we observed in LUAD would remain true in COAD. Therefore, we generated CRISPR/Cas9- mediated knockout of *KSR1* in four COAD cell lines with varying *PIK3CA* status: SW480 (*KRAS^G12C^/PIK3CA^WT^*) LoVo (*KRAS^G13D^/PIK3CA^WT^*), LS174T (*KRAS^G12D^/PIK3CA^mut^*), and T84 (*KRAS^G13D^/PIK3CA^mut^*). *In vitro* ELDAs performed with *KSR1* KO in the four COAD cell lines demonstrated a 2-3-fold significant decrease in SIC-frequency compared to NT cells. Further, *KSR1* KO prevented the trametinib-induced increase in SICs in the four COAD cell lines (Fig. 4 A-B), demonstrating that the KSR1 effect on SICs in COAD is independent of *PIK3CA* mutational status. Further, treatment with BI-3406 in NT cells prevented trametinib-induced SIC increase in the cell lines with wildtype *PI3KCA* status (SW480 and LoVo), but not in *PI3KCA^mut^* cell lines (LS174T and T84), consistent with our LUAD findings (Fig. 2 E). In *KSR1* KO cells, combination of trametinib with BI-3406 did not further affect SIC frequency, concordant with SOS1 acting upstream of KSR1 in the RTK/RAS pathway (*SI Appendix, Fig. S13*).

**Fig. 4.**
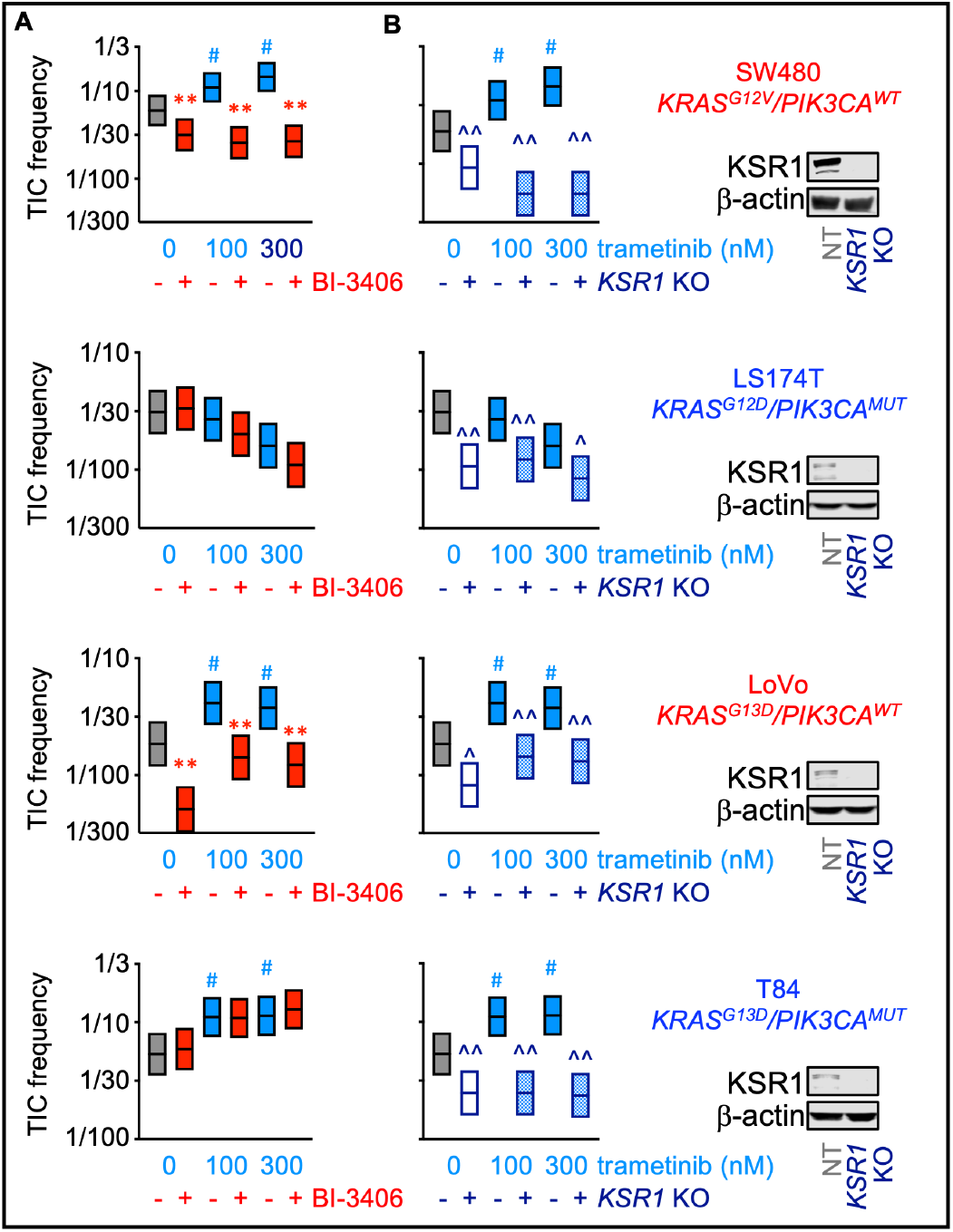
*KSR1* KO and SOS1 inhibition show differential inhibition of basal and trametinib-induced SICs in *KRAS*-mutated COAD cells. *(A)* SIC frequency from *in situ* ELDAs in the indicated COAD cell lines pre-treated with trametinib for 72 hours to upregulate SICs, and then left untreated or treated with the SOS1 inhibitor BI-3406. The *KRAS* and *PIK3CA* mutational status for each cell line is indicated. *(B)* SIC frequency from *in situ* ELDAs in the indicated NT and *KSR1* KO COAD cells pre-treated with trametinib for 72 hours. Western blots of WCLs for KSR1 and β-actin are shown on the right. # p < 0.05 vs untreated; ## p<0.01 vs. untreated for SIC upregulation by MEK inhibitor treatment vs. untreated controls. ** p<0.01 for SIC inhibition by BI-3406 treatment compared to untreated controls. x^ p<0.05; ^^ p<0.01 for *KSR1* KO compared to untreated controls. Data are representative of three independent experiments.

### KSR1 regulation of SIC survival in *KRAS*-mutated COAD is dependent on interaction with ERK

KSR1 mediates ERK-dependent signaling in transformed and untransformed cells via direct interaction between its DEF domain and ERK (40, 63, 64). A KSR1 transgene deficient in binding ERK due to engineered mutation in the DEF- domain, KSR1^AAAP^ (40), was expressed in *KSR1* KO colorectal adenocarcinoma cell line HCT116 (Fig. 5 A), which rescued decreased MEK but not ERK phosphorylation in *KSR1* KO cells (*SI Appendix, Fig. S14*). Similar to H460 cells, HCT116 cells demonstrate a high (>50%) SIC frequency by *in situ* ELDA, therefore single cell clonogenicity, colony formation in soft agar, and ALDH activity were assessed as *in vitro* measures of SICs. Expression of KSR1^AAAP^ in *KSR1* KO cells failed to rescue ALDH activity, single cell clonogenicity, or anchorage-independent growth by soft agar assay to the level observed with wildtype KSR1 addback, demonstrating the necessity of ERK interaction on KSR1 regulation of SICs (Fig. 5 B-D). To assess KSR1 function in a preclinical setting, an *in vivo* limiting dilution analysis was performed. Notably, a 70-fold decrease in the proportion of TICs was found in the *KSR1* KO cells compared to those with NT cells, demonstrating the significant impact of KSR1 on TICs in COAD (Fig. 5 E). A more extensive single cell clonogenicity assay assessing growth in the HCT116 cells treated with 100 or 300 nM of trametinib further demonstrated that only in those cells expressing an intact KSR1 protein did trametinib enhance clonogenicity, while *KSR1* KO and KSR1^AAAP^ cells that lack the ability to bind ERK were insensitive to SIC induction by trametinib (Fig. 5 F).

**Fig. 5.**
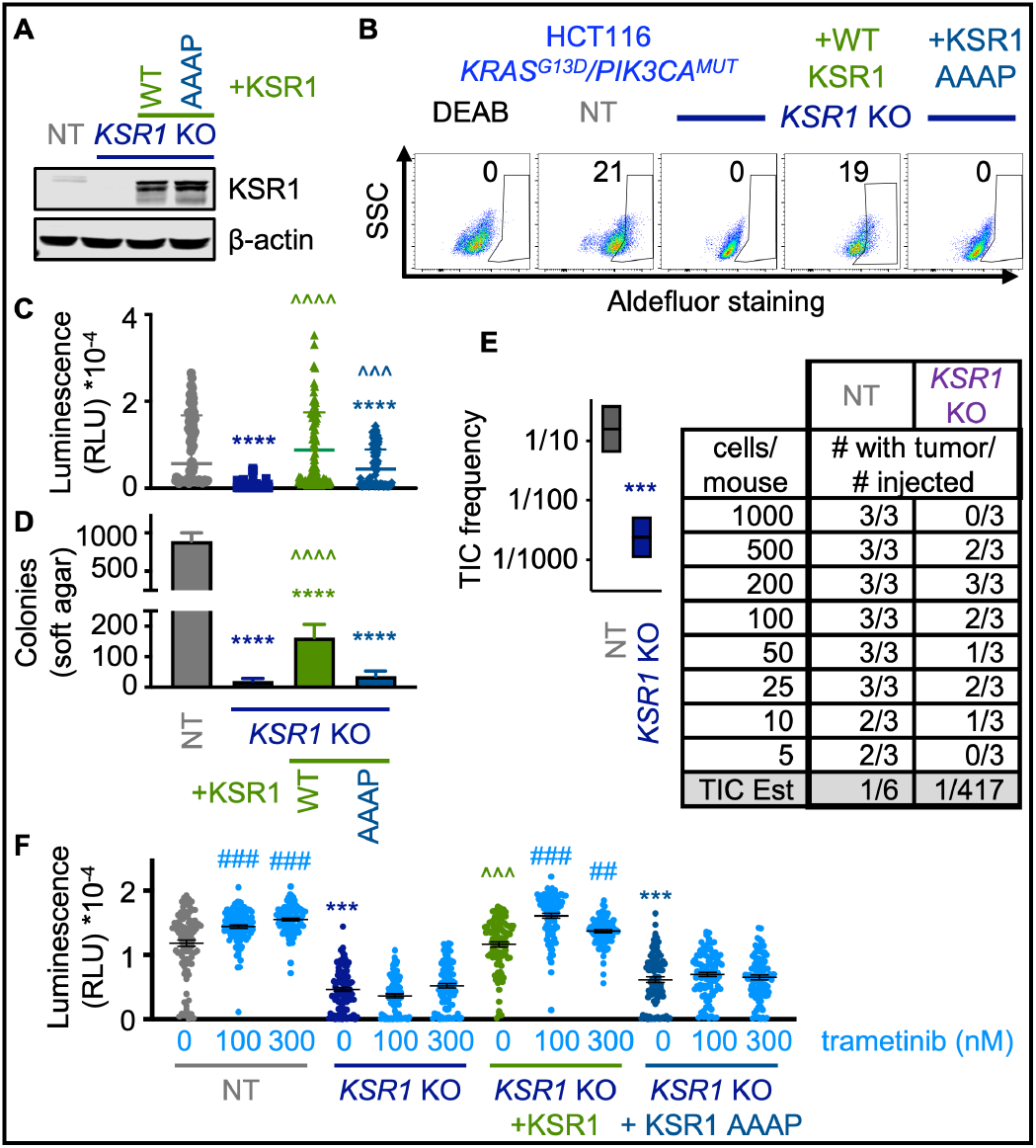
KSR1 regulation of TICs/SICs in COAD is dependent on interaction with ERK and relevant *in vivo*. *(A)* Western blot for KSR1 and β-actin loading controls from WCLs of HCT116 (*KRAS^G13D^/PIK3CA^mut^*NT, *KSR1* KO, *KSR1* KO + *KSR1* addback, and *KSR1* KO+ERK-binding mutant KSR1 (KSR1^AAAP^) addback cells. *(B)* Aldefluor staining for ALDH enzyme activity in the indicated cells including a DEAB negative control. *(C)* Single cell colony forming assays. Cells were single cell plated in non-adherent conditions, and colony formation was scored at 14 days by CellTitre Glo. Each individual point represents a colony. *(D)* Soft agar colony forming assay. 1×10^3^ cells per well were plated in 0.4% agar, and colony formation was scored at 14 days. *(E) In vivo* limiting dilution analysis data showing frequency of TICs in Non-targeting Control (NT) and KSR1 KO HCT116 COAD cells. The indicated numbers of cells were injected into the shoulder and flank of NCG mice (Charles River). Tumors were scored at 30 days. *(F)* Single cell colony forming assay in H460 cells pre-treated with the indicated doses of trametinib for 72 h. Cells were single cell plated in non-adherent conditions, and colony formation was scored at 14 days by CellTitre Glo. Each individual point represents a colony. ## p<0.01, ### p<0.001 vs. untreated for SIC upregulation by MEK inhibitor treatment vs. untreated controls; *** p<0.001, ****p<0.0001 for TIC/SIC inhibition by *KSR1* KO versus controls; ^^^ p<0.001, ^^^^ p<0.0001 versus *KSR1* KO.

### SOS1 and KSR1 disruption prevent trametinib resistance in *KRAS*-mutated cells

To assess the effect of SOS1 and KSR1 disruption on outgrowth of trametinib-resistant cells, we utilized multi-well *in situ* resistance assays (65) in which cells are grown on a 96-well plate and treated with trametinib alone or in combination with BI-3406. Wells are scored twice weekly to assess for wells with >50% confluency to determine the presence of resistance. Of the five LUAD cell lines tested, SOS1 inhibition with BI-3406 prevented outgrowth of trametinib-resistant cells in (*KRAS^G12V^/PI3KCA^WT^*) and H358 cells (*KRAS^G12C^/PI3KCA^WT^*), but not in LUAD cell lines with either a *PIK3CA* co-mutation (LU99A), *KRAS^Q61^*mutation (Calu6), or both (H460) (Fig. 6A-E). In contrast, *KSR1* KO was able to prevent outgrowth of trametinib-resistant cells in H460 LUAD cells (Fig. 6 F) and in the HCT116 COAD cell line (*KRAS^G13D^/PIK3CA^mut^*) (Fig. 6 G). To determine whether interaction with ERK was necessary for the KSR1 effect on trametinib resistance, we further tested whether expression of ERK-binding mutant KSR1^AAAP^ in *KSR1* KO cells could rescue trametinib-resistant outgrowth. KSR1^AAAP^ partially restored outgrowth relative to *KSR1* KO cells while wildtype KSR1 completely restored outgrowth (Fig. 6 F), suggesting KSR1 interaction with ERK affects trametinib resistance but may be occurring in combination with other KSR1-dependent effects.

**Fig. 6.**
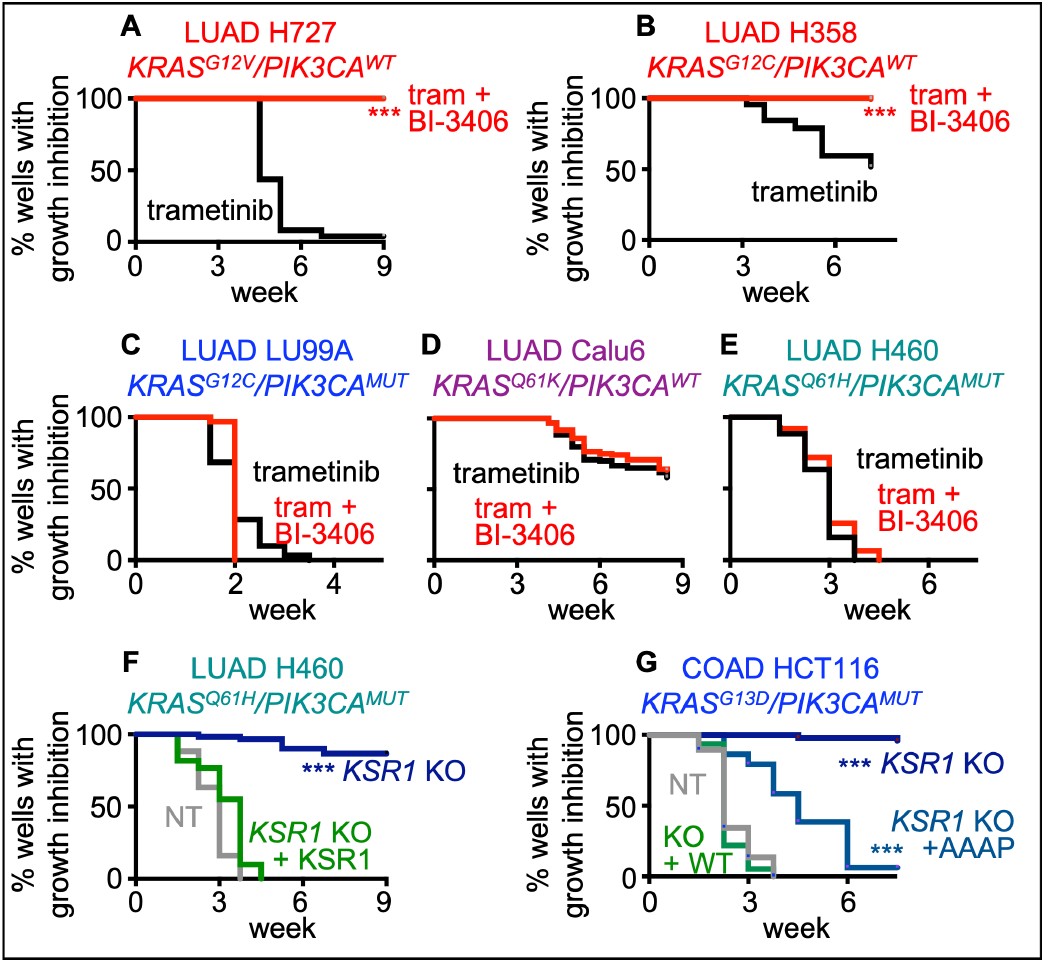
SOS1 inhibition and *KSR1* KO delay outgrowth of trametinib-resistant cells in multi-well resistance assays depending upon the KRAS mutational status. Multi-well resistance assays were performed as outlined in the Materials and Methods. *(A-E)* Trametinib resistance in *KRAS^G12X^*/*PIK3CA*^WT^ H727 *(A)* and H358 *(B)*, *KRAS^G12C^*/*PIK3CA^MUT^*LU99A *(C)*, *KRAS^Q61X^*/*PIK3CA^WT^* Calu6 cells treated with trametinib *(D)*, or *KRAS^Q61X^*/*PIK3CA^MUT^* H460 cells *(E)* treated with an EC_85_ dose of trametinib with and without SOS1 inhibitor BI-3406. *(F-G)* Trametinib resistance in control and *KSR1* KO *KRAS^Q61H^*-mutated/*PIK3CA^MUT^*H460 LUAD cells *(F)* and *KRAS^G13D^*-mutated/*PIK3CA^MUT^*HCT116 COAD cells *(G)*. In *(G)*, rescue of *KSR1* KO using either WT KSR1 or a KSR1^AAAP^ ERK-binding mutant was also tested for its effect on trametinib resistance. Data from *N*=3 independent experiments were combined to generate Kaplan-Meier curves. *** p<0.001 vs single drug treatment *(A-E)* or NT controls *(F-G)*.

## Discussion

Within the RTK/RAS pathway, there is a hierarchical dependency of downstream signaling pathways depending upon the specific RAS mutation, with KRAS predominantly signaling downstream to the RAF/MEK/ERK pathway (9, 66–69). Thus, targeting MEK is an attractive option for treating patients with *KRAS*-mutated tumors. Unfortunately, trametinib monotherapy is largely ineffective due both to the loss of ERK-dependent negative feedback control of RTKs (adaptive resistance (5, 14–19, 21, 22)) as well as subsequent selection of tumor initiating cells through therapeutic-pressure over-time (acquired resistance (5, 13, 21, 23)). Previous studies designed to identify MEK inhibitor co-targets have identified combinations that can overcome adaptive resistance (22, 32, 70, 71), but have not examined the extent to which these combinations may prevent the acquisition of acquired resistance. Here, we provide an experimental framework for evaluating both adaptive and acquired resistance to RTK/RAS pathway targeted therapies and use this framework to show that, with knowledge of the specific mutational profile (i.e. specific *KRAS* mutation and *PIK3CA* co-mutation status), vertical inhibition of RTK/RAS signaling can enhance the overall effectiveness of MEK inhibitors in *KRAS*-mutated cancer cells.

Essential to building this framework is having reliable experimental approaches that model each step of the evolution of a cancer cell due to therapeutic pressure and then to use this framework when assessing novel drug combinations. The ideal drug combination would (i) enhance the efficacy of an oncogene-targeted therapy to overcome intrinsic/adaptive resistance, (ii) limit the survival of TICs, which are the subset of drug-tolerant persister cells capable of driving adaptive resistance (47, 49, 57, 72), and (iii) delay the onset of and/or block the development of resistant cultures. To examine the extent to which combination therapies enhance the efficacy of an oncogene-targeted therapy to overcome intrinsic/adaptive resistance in *KRAS*-mutated cancers, we assessed drug-drug synergy in 3D spheroid cultures (Fig. 1). 3D culture conditions are essential to the assessment of drug-drug synergy in RTK/RAS-mutated cancers. *KRAS*-mutated cell lines originally classified as KRAS-independent in 2D adherent culture (73–77) require KRAS expression (78–81) or become sensitized to KRAS^G12C^ inhibitors (82) in 3D culture conditions. Further, we and others have shown that inhibition or deletion of proximal RTK signaling intermediates SOS1 (31, 32, 83), SOS2 (18, 69), and SHP2 (22, 83–85) inhibit proliferation of RTK/RAS mutated cancers and synergize with therapies targeting the RTK/RAS pathway, but only under 3D culture conditions. To assess enrichment of SICs within the therapy-tolerant persister cell population and the extent to which combination therapies can block this enrichment, we perform ELDAs (9, 86) in 3D culture conditions (Figs. 2-4) that allow us to estimate the frequency of SICs within a cell population and show increased SIC frequency when *KRAS*-mutated cells are pre-treated with trametinib. This enrichment of SICs upon trametinib treatment confirms that beyond adaptive resistance, there is likely underlying molecular heterogeneity in *KRAS*-mutated cancers associated with drug-tolerant persister (DTP) cells that allow for acquired resistance to trametinib over time. To assess the extent to which therapeutic combinations limit the development of acquired resistance, we use *in situ* resistance assays that our laboratory developed as a hybrid approach that combines elements of time-to-progression assays (71, 87) and cell outgrowth assays originally describe by the Jänne laboratory (88, 89). These longitudinal studies of acquired resistance act as a cell-culture surrogate of multi-individual trials that should be performed prior to testing therapeutic combinations *in vivo* (65).

Using this framework, we found SOS1 inhibition using BI-3406 both enhanced the efficacy of trametinib by preventing reactivation of AKT and ERK signaling and prolonged the therapeutic window of trametinib by targeting SICs and thereby preventing the development of acquired resistance in *KRAS^G12/G13^*-mutated LUAD and COAD cells. However, the effectiveness of BI-3406 was lost either in *KRAS^Q61^*-mutated cells or in cells harboring *PIK3CA* co-mutations. For *KRAS*-mutated cells harboring *PIK3CA* co-mutations, constitutive PI3K-AKT signaling bypasses the RTK-dependent PI3K activation that normally occurs due to loss of ERK-dependent negative feedback after trametinib treatment, thereby abrogating the ability of proximal RTK pathway inhibitors including SOS1 to synergizes with trametinib. These data are further consistent with previous studies showing that SOS2 was required for mutant KRAS-driven transformation, but that transformation could be restored in *Sos2* KO cells by expression of activated PI3K (18).

In *KRAS^Q61^*-mutated cells, the inability of SOS1 inhibitors to synergize with trametinib is likely due to the heterogeneous molecular behavior of codon-specific KRAS mutations with regard to GTP/GDP cycling (Fig. 1G, S5) (90); while G12, G13, and Q61 mutants all show reduced GAP-dependent GTP hydrolysis, Q61 mutants that show dramatically reduced intrinsic GTP-hydrolysis compared to G12/G13. The extremely low level of GTP hydrolysis (KRAS inactivation) seen in Q61 mutants makes them much less dependent on RASGEFs for their continued activation compared to G12/G13 mutants (36, 37). Indeed, others have shown that SHP2 and SOS1 inhibitors enhance the killing effects of MEK inhibitors in *KRAS^G12X^*- and *KRAS^G13X^*-mutated, but not *KRAS^Q61X^*- mutated, tumors (22, 32). Since the ineffectiveness of MEK inhibitors has been attributed not only to feedback RTK-PI3K signaling but also to compensatory ERK reactivation (5, 19, 20), we asked whether deletion of the RAF/MEK/ERK scaffold KSR1 could cause deep ERK inhibition and enhance the effectiveness of trametinib in *KRAS*-mutated cancer cells that were insensitive to SOS1 inhibition.

We found that in *KRAS^Q61^*/*PIK3CA* mutated LUAD cells, which would be the least sensitive to SOS1 inhibition, *KSR1* KO synergized with trametinib to inhibit ERK signaling, thereby limiting survival and significantly decreasing TIC frequency *in vivo*. Although *PIK3CA* co-mutations are rare in *KRAS*-mutated LUAD, they commonly occur in COAD (91, 92). Thus, we shifted our assessment of *KSR1* KO to COAD cells, where we found *KSR1* KO inhibited the trametinib-induced enrichment of SICs in *KRAS*- mutated COAD cells regardless of *PIK3CA* mutation status. We further showed that these effects were due to KSR1 scaffolding function, as an ERK-binding mutant (KSR1^AAAP^) failed to rescue SIC properties (ALDH activity, soft agar growth, clonogenicity) compared to a WT KSR1 transgene.

This finding is consistent with KSR1 function as a RAF/MEK/ERK scaffold and with our previous studies showing KSR1-ERK signaling is essential to mutant *RAS*- driven transformation (38, 40, 93–95). Here, we extend our understanding of KSR1 scaffolding to show that it is essential to the therapeutic response. KSR1 disruption enhances trametinib-induced destabilization of the MEK-ERK complex approximately 30-fold, increasing the sensitivity to the MEK inhibition. These findings, when coupled to our previous data showing that PI3K/AKT signaling is independent of KSR1 (38, 40) and KSR1 depletion inhibited transformation in *KRAS*/*PIK3CA* co-mutated COAD cells (93–95), give further support to compensatory ERK reactivation as a key component of adaptive resistance to trametinib that can be inhibited by targeting KSR1. Further, the finding that *ksr1*^-/-^ mice are phenotypically normal but resistant to cancer formation (41, 42) highlights the potential of targeting KSR1 to achieve a high therapeutic index. A recently developed KSR inhibitor increased the potency of MEK inhibitors, demonstrating that the use of KSR and MEK inhibitors may be a promising combination therapeutic strategy (59).

In addition to overcoming intrinsic/adaptive resistance, optimal combination therapies would also delay the development of acquired resistance and prolong the window of efficacy for trametinib treatment. Unfortunately, most studies of resistance to RTK/RAS pathway inhibitors including trametinib focus either on synthetic lethality during a short treatment window (0-28 days) (17, 22, 23, 70, 96) or study resistance in a few cell lines established by dose-escalation over several months (97) rather than determining the extent to which combination therapies can delay the onset of acquired resistance. Using *in situ* resistance assays to assess acquired resistance to RTK/RAS pathway inhibitors in large cohorts of cell populations (65), we found that SOS1 inhibition inhibited the development of trametinib resistance in *KRAS^G12^*-mutated LUAD cells, which represent the majority of *KRAS*-mutated LUADs. Mutations in RTK/RAS pathway members, including KRAS, occur in 75-90% of LUAD, and RTK pathway activation is a major mechanism of acquired resistance in LUADs with *EGFR* mutations (98–107), mutations in alternative RTKs (108–117)), or *KRAS^G12^* (118–120)) or non-G12C (14–18) mutations likely due to RTK/RAS pathway addiction in these tumors (109, 121–124). In addition to SOS1, the RASGEF SOS2 and the phosphatase/adaptor SHP2 represent proximal RTK signaling intermediates and potential therapeutic targets whose inhibition may limit resistance to RTK/RAS pathway inhibitors in LUAD. In parallel studies, we found that inhibiting proximal RTK signaling by either SHP2 inhibition (65) or *SOS2* deletion (125) delayed or inhibited the development of osimertinib resistance in *EGFR*-mutated LUAD cells. Based on these data, we propose that proximal RTK inhibition as a therapeutic strategy to delay resistance to RTK/RAS pathway targeted therapies in a majority of LUADs. However, SOS1 inhibition failed to inhibit resistance in cells with either *KRAS^Q61^* mutations or with co-occurring *PIK3CA* mutations. In these settings, we found that *KSR1* KO significantly reduced the number of trametinib-resistant colonies suggesting that targeting KSR1 may be a better approach in these genetic backgrounds. While co-occurring *KRAS* and *PIK3CA* mutations are rare in LUAD, ∼ 1/3 of *KRAS*- mutated colorectal cancers harbor *PIK3CA* mutations. Thus, we propose that KSR1 may be a better co-therapeutic target compared to SOS1 in COAD.

Our study provides a framework for evaluating and selecting optimal combination therapies to limit both intrinsic/adaptive and acquired resistance to RTK/RAS pathway targeted therapies. Using this framework, we demonstrated that either SOS1 inhibition or KSR1 disruption can increase the efficacy of trametinib and prevent both intrinsic and acquired resistance with genotype-specificity; SOS1 inhibition was more effective in cells harboring *KRAS^G12/G13^* mutations with wild-type *PIK3CA*, whereas *KSR1* KO was more effective in cells with co-occurring *PIK3CA* mutations. While strategies to inhibit KSR1 are still under development (59, 126), SOS1 inhibitors BI 1701963 [NCT04111458; NCT04975256] and MRTX0902 [NCT05578092] are currently being evaluated in Phase1/2 studies for treatment of *KRAS*-mutated cancers either alone or in combination with trametinib or the KRAS^G12C^ inhibitor adagrasib. Our finding that SOS1 inhibitors delay resistance to trametinib only in *KRAS^G12/G13^*-mutated cells that lack *PIK3CA* co-mutations has implications for understanding which patient populations will likely benefit from combined SOS1/MEK inhibition and should inform future clinical trial design for SOS1 inhibitor combinations.

## Materials and Methods

### Cell culture

Lung and colon cancer cell lines were purchased from ATCC or JCRB (LU99A). After receiving the cells, they were expanded and frozen at passage 3 and 4; cell were passaged once they became 70-80% confluent and maintained in culture for 2-3 months before thawing a new vial as prolonged passaging can alter TIC frequency (127). Cell lines were cultured at 37°C and 5% CO2. Cells were passaged in either RPMI [H727, A549, H358, LU99A, H460] or DMEM [Calu6, H650, H1155, SW620, SW480, LS174T, LoVo, T84, HCT116] supplemented with 10% FBS and 1% penicillin/streptomycin. For signaling experiments, cells were seeded in 24-well micropatterned AggreWell 400 low-attachment plates (StemCell) at 9 x 10^5^ cells/ well in 2 mL of medium. 24-h post plating, 1 mL of media was replaced with 2x inhibitor. Cells were treated with inhibitor for 0 – 72 hours; for all treatments >24-h, half of the media was removed and replaced with fresh 1× inhibitor daily.

### Production of recombinant lentiviruses and sgRNA studies

Lentiviruses were produced by co-transfecting MISSION lentiviral packaging mix (Sigma) into 293T cells using Mirus TransIT-Lenti transfection reagent (Mirus Bio # MIR6605) in Opti-MEM (Thermo Scientific #31-985-062). 48-h post-transfection, viral supernatants were collected and filtered. Viral supernatants were then either stored at −80°C or used immediately to infect cells in combination with polybrene at 8 μg/mL.

### Generation of pooled genomic *SOS1* KO cell lines

For *SOS1* KO studies, cells were infected with lentiviruses based on pLentiCRISPRv2 with either a non-targeting sgRNA (NT) or a sgRNA targeting SOS1 (83). 48-h post-infection, cells were selected in 4 μg/mL Puromycin (Invitrogen). 7-10 days after selection, cells were analyzed for SOS1 or KSR1 expression. Cell populations showing >80% deletion were used for further study.

### Generation of cells expressing PI3K or ERK2-MEK1 proteins

For p110α expression experiments, cells were infected with lentiviruses expressing WT p110α (Addgene #116771), p110α^H1047R^ (Addgene #116500), or pHAGE-puro empty vector (Addgene #118692) (128, 129). 48-h post-infection, cells were selected in 4 μg/mL Puromycin (Invitrogen). For ERK2-MEK1 fusion experiments, cells were infected with retroviruses expressing WT ERK2-MEK1, nuclear localized ERK2-MEK1 LA, or YFP empty vector (62). YFP+ cells were selected by fluorescence-activated cell sorting. Four days after selection or sorting, cells were analyzed for p110α or ERK2-MEK1 fusion protein expression.

### Generation of clonal genomic *KSR1* KO cell lines

sgRNA sequences targeting KSR1 or non-targeting control were inserted into pCAG-SpCas9-GFP-U6-gRNA (Addgene #79144, gift of Jizhong Zou). PEI transfection was used to insert pCAG-SpCas9-GFP-U6-sgKSR1 or non-targeting control into H460 and HCT116 cells. GFP-selection by fluorescence-activated cell sorting was performed 48-h post-transfection, and colonies were grown out with the use of cloning rings.

### Generation of pooled genomic *KSR1* KO cell lines

sgRNA sequences targeting KSR1 or non-targeting control were inserted into pLentiCRISPRv2GFP (Addgene #82416). The constructs were PEI transfected into HEK293T cells along with psPAX2 lentiviral packaging construct (Addgene #12259) and pMD2.G envelope construct (Addgene #12259). Lentivirus-containing media was harvested at 96-h, and added to SW480, LoVo, LS174T, and T84 cells. GFP+ cells were selected by fluorescence-activated cell sorting.

### Generation of KSR1 addback in clonal genomic *KSR1* KO cell lines

Murine KSR1 was cloned into MSCV-IRES-KSR1-GFP (Addgene #25973) and PEI transfected into HEK293T cells along with pUMVC retroviral packaging construct (Addgene # 8449) and VSVG envelope construct (Addgene #8454). Lentivirus-containing media was harvested at 96-h and added to clonal *KSR1* KO HCT116 and H460 cells. GFP-selection was performed by fluorescence-activated cell sorting 48-h post-transfection.

### Single Cell Colony-forming Assay

Cells were DAPI-stained for viability determination and live cells were single cell sorted as one cell per well into a 96-well, U-bottom, non-adherent plate (Takara Bio). Cells were grown for 14 days, after which colony formation was determined using CellTiter Glo® viability assay (Promega) performed according to manufacturer instructions.

### Flow cytometry

Cells were plated in 10 or 6 cm tissue-culture treated plates and allowed to adhere for 24-48-h prior to drug treatment. Once cells were 50-75% confluent, cells were treated with the indicated concentration of trametinib or selumetinib for 72-h. After the 72-h treatment, cells were harvested by trypsinization, spun down, resuspended in Aldefluor Assay Buffer (StemCell) at 1 × 10^6^ cells/mL, and stained for ALDH activity using the Aldefluor Assay Kit per manufacturer’s instructions. An aliquot of cells was pre-treated with the ALDH inhibitor DEAB, which inhibits ALDH enzymatic activity and is thus used as a negative gating control. Data were analyzed using FloJo with and are presented as the % of cells showing ALDH activity over DEAB controls.

### Soft Agar Colony-forming Assay

1 x 10^3^ cells per dish were plated onto 35mm dishes in 0.4% NuSieve Agarose (Lonza #50081). Six replicates of each condition were plated. At 14 days, colony formation was assessed by counting colonies > 100 μM in their largest diameter.

### Cell lysis and Western blotting

Cells were lysed in RIPA buffer (1% NP-40, 0.1% SDS, 0.1% Na-deoxycholate, 10% glycerol, 0.137 M NaCl, 20 mM Tris pH [8.0], protease (Biotool #B14002) and phosphatase (Biotool #B15002) inhibitor cocktails) for 20 min at 4°C and spun at 10,000 RPM for 10 min. Clarified lysates were boiled in SDS sample buffer containing 100 mM DTT for 10 min prior to western blotting. Proteins were resolved by sodium dodecyl sulfate-polyacrylamide (Criterion TGX precast) gel electrophoresis and transferred to nitrocellulose membranes. Western blots were developed by multiplex Western blots were developed by multiplex Western blotting using anti-SOS1 (Santa Cruz sc-256; 1:500), anti-β-actin (Sigma AC-15; 1:5,000 or Santa Cruz Biotechnology sc-47778, 1:2000 dilution), anti-HSP90 (Santa Cruz sc-515081; 1:500), anti-α-tubulin (Abcam ab89984; 1:2000); anti-pERK1/2 (Cell Signaling 4370; 1:1,000), anti-ERK1/2 (Cell Signaling 4696; 1:1000), anti-pAKT Ser473 (Cell Signaling 4060; 1:1000), anti-AKT (Cell Signaling 2920; 1:1000), anti-KSR1 (Abcam ab68483, 1:750 dilution), anti-pMEK1/2 (Cell Signaling 9154; 1:1000), anti-MEK1/2 (Cell Signaling 4694; 1:1000), anti-pRSK (Cell Signaling 9344; 1:1000), anti-RSK (B-4) (Santa Cruz sc-74575; 1:1000) anti-pS6 Ser235/236 (Cell Signaling 4858; 1:1000), anti-S6 (Cell Signaling 2317; 1:1000) primary antibodies. Western blot protein bands were detected and quantified using the Odyssey system (LI-COR).

### G-LISA

GTP-bound RAS levels were assessed with a RAS G-LISA Assay Kit per manufacturer’s instructions (Cytoskeleton INC., #BK131). 9 × 10^5^ cells were plated in a 3D 24-well plate. Media was changed every 24 hours. Two replicates were plated per both parental and SOS 1 KO cell line. Once treatment was complete, cells were collected by washing the well with media and collecting all media. Two PBS washes were done before the Cell Lysis Buffer from the G-LISA kit was used per protocol specifications. To assist with the lysis, cells were flash frozen and stored at −80°C prior to continuing with the protein lysis collection. A protein assay was completed immediately after cells were thawed and centrifuged. After samples were equalized and sufficient lysate mixed with equal parts binding buffer, 50 μL were added to the wells; a negative and positive well were also used for the experiment. The plate was kept cold (4°C) and on an orbital microplate shaker at 400 rpm for 30 min before continuing with anti-RAS primary antibody incubation at 1:100 dilution for 30 minutes at room temperature followed by a wash cycle and continuation of the protocol per manufactures instructions.

### Extreme Limiting Dilution Analysis (ELDA)

#### In vivo

In equal volumes of 50% Cultrex® Basement Membrane Extract (R&D Systems) 50% Dulbecco’s Modified Eagle Medium (Cytiva), dilutions ranging from 5 to 1000 cells were injected subcutaneously into the shoulder and hip of 6-8-week-old triple immunodeficient male NOD-Prkdc^em26Cd52^Il2rg^em26Cd22^/NjuCr (NCG, Charles River) mice. Mice were monitored for tumor formation by palpation. Once tumor size reached 1cm^3^, mice were sacrificed and tumors were excised. TIC frequency and significance between groups was calculated by ELDA website https://bioinf.wehi.edu.au/software/elda/ (86).

#### In situ

Cells were seeded in 96-well ultra-low attachment flat bottomed plates (Corning Corstar #3474) at decreasing cell concentrations (1000 cells/well – 1 cell/well) at half log intervals (1000, 300, 100,30, 10, 3, 1), 12 wells per condition with the exception of the 10 cells/well condition, for which 24 wells were seeded. Cells were cultured for 7-10 days, and wells with spheroids >100 mM were scored as spheroid positive. TIC frequency and significance between groups was calculated by ELDA website https://bioinf.wehi.edu.au/software/elda/ (86). To assess the effect of trametinib on TIC frequency, cells were left untreated or were pre-treated with the indicated dose of trametinib or selumetinib for 72-h, after which cells were rested for 48-72-h prior to plating. To assess the effect of SOS1 inhibition, cells were seeded ± 300 nM BI-3406.

### Proximity Ligation Assay

Cells were seeded at a density of 1 × 10^4^ cells per well in (16-well Nunc Lab-Tek Chamber Slides #178599PK, ThermoFisher) in 200 µl of normal media. Once cells were adhered to the plate, 200 µl of normal media or drugged media was added for a final volume of 400 µL and allowed to rest for 24 h. Media was aspirated from the wells followed by a single cold PBS wash. Cells were then fixed with 4% formaldehyde for 20 minutes followed by permeabilization with 0.5% Triton X-100 solution with four 5 min wash cycles (200 µL per wash). After blocking, cells were incubated with rabbit anti-ERK (1:250; #4695) and mouse anti-MEK (1:50; #4694) primary antibodies overnight at 4°C. The following day, the primary antibodies were slowly aspirated followed by wash buffer A (#DUO82049) and incubation with PLA probes for Rabbit/Mouse (#DUO92102) per manufacturer’s instructions. Ligation and amplification were performed per the provided protocol (#DUO92008). Final washes were performed with wash buffer B before mounting the slides with the *in situ* mounting medium with DAPI (#DUO82040) and imaged at 60x. Images were normalized to background staining of NT wells incubated with anti-ERK or anti-MEK antibodies alone and analyzed using Fiji and ImageJ. Particle size was thresholded at 0.1 µM^2^, and the number and size of particles per cell were quantified. Per condition, 6 – 10 cells were analyzed from three independent wells (>20 cells per condition).

### Bliss Independence Analysis for Synergy

Cells were seeded at 750 cells per well in 100 mL in the inner-60 wells of 96-well ultra-low attachment round bottomed plates (Corning #7007) or Nunc NucleonSphera microplates (ThermoFisher # 174929) and allowed to coalesce as spheroids for 24-48 h prior to drug treatment. Cells were treated with drug for 96 h prior to assessment of cell viability using CellTiter Glo® 2.0 (Promega). For all studies, outer wells (rows A and H, columns 1 and 12) were filled with 200 mL of PBS to buffer inner cells from temperature and humidity fluctuations.

Triplicate wells of cells were then treated with increasing concentrations trametinib alone, BI-3406 alone, or the combination of trametinib + BI-3406 in a 9 × 9 matrix of drug combinations on a similog scale for 72 h (adherent cultures) or 96 h (spheroids). Cell viability was assessed using CellTiter Glo® 2.0 (30 mL/well). Luminescence was assessed using a Bio-Tek Cytation five multi-mode plate reader. Data were normalized to the maximum luminescence reading of untreated cells, and individual drug EC_50_ values were calculated using Prism9 by non-linear regression using log(inhibitor) vs. response. For all drug-treatment studies, the untreated sample for each cell line was set to 100%. This would mask any differences in 3D cell proliferation seen between cell lines. Excess over Bliss was calculated as the Actual Effect – Expected Effect as outlined in (83). The SUM EOB is calculated by taking the sum of excess over bliss values across the 9 × 9 treatment matrix. EOB values > 0 indicate increasing synergy.

### Resistance Assays

Cells were plated at low density (250 cells/well) in replicate 96-well plates, and each plate was treated with the indicated doses of trametinib ± BI-3406.

Wells were fed and assessed weekly for outgrowth, wells that were > 50% confluent were scored as resistant to the given dose of trametinib. Data are plotted as a Kaplan-Meier survival curve; significance was assessed by comparing Kaplan-Meyer Meier curves using Prism 9.

## List of Key Resources

KSR1 antibody: Abcam ab68483, 1:750 dilution

β-actin antibody: Santa Cruz Biotechnology sc-47778, 1:2000 dilution

β-actin antibody: Sigma AC-15; 1:5,000 dilution

HSP90 antibody: Santa Cruz sc-515081; 1:500 dilution

SOS1 antibody: Santa Cruz sc-256; 1:500 dilution

a-tubulin antibody: Abcam ab89984 1:2000 dilution

pERK1/2 antibody: Cell Signaling 4370; 1:1,000 dilution

ERK1/2 antibody: Cell Signaling 4696; 1:1000 dilution

ERK 1/2 antibody: Cell Signaling 4695; 1:200 dilution (proximity ligation assay)

pAKT Ser473 antibody: Cell Signaling 4060; 1:1000 dilution

AKT antibody: Cell Signaling 2920; 1:1000 dilution

pMEK 1/2 antibody: Cell Signaling 9154; 1:1000 dilution

MEK 1/2 antibody: Cell Signaling 4694; 1:1000 (western)/1:50 (proximity ligation assay)

pRSK antibody: Cell Signaling 9344; 1:1000 dilution

RSK (B-4) antibody: Santa Cruz sc-74575; 1:1000 dilution

pS6 Ser235/236 antibody: Cell Signaling 4858; 1:1000 dilution

S6 antibody: Cell Signaling 2317; 1:1000 dilution

KSR1 sgRNA sequences:

CR1 5’ TTGGATGCGCGGCGGGAAAG 3’

CR2 5’ CTGACACGGAGATGGAGCGT 3’

NT sgRNA sequence: 5’ CCATATCGGGGCGAGACATG 3’

SOS1 sgRNA sequence: 5’ GCATCCTTTCCAGTGTACTC 3’

Plasmid catalog numbers listed in the sections above.

## Acknowledgments

We thank the UNMC Cell Analysis Facility and UNMC Animal Facility. We thank Gordon Mills (MD Anderson Cancer Center, Houston, TX) and Christopher Vakoc (Cold Spring Harbor Laboratory, Cold Spring Harbor, NY) and Melanie Cobb (UT Southwestern, Houston, TX) for plasmids.

