## Supplemental Figures S1 - S14 for "SOS1 and KSR1 modulate MEK inhibitor responsiveness to target resistant cell populations based on PI3K and KRAS mutation status"

Brianna R. Daley[1,2], Heidi M. Vieira[1,3], Chaitra Rao[3,4], Jacob M. Hughes[2], Zaria M. Beckley[2], Dianna H. Huisman[3], Deepan Chatterjee[5], Nancy E. Sealover[2], Katherine Cox[2], James W. Askew[3], Robert A. Svoboda[6], Kurt W. Fisher[6], Robert E. Lewis[3*], and Robert L. Kortum[2*]
[1]These authors contributed equally to this paper
[2]Department of Pharmacology and Molecular Therapeutics, Uniformed Services University of the Health Sciences, Bethesda, MD
[3]Eppley Institute for Research in Cancer and Allied Diseases, University of Nebraska Medical Center, Omaha, NE
[4]Current address: Wells Center for Pediatric Research, Indiana University School of Medicine, Indianapolis, IN
[5]Integrative Physiology and Molecular Medicine, University of Nebraska Medical Center, Omaha, NE
[6]Department of Pathology and Microbiology, University of Nebraska Medical Center, Omaha, NE
* Corresponding Authors

Robert E. Lewis
Eppley Institute for Research in Cancer and Allied Diseases
University of Nebraska Medical Center
Omaha, NE 68198

Robert L. Kortum
Uniformed Services University of the Health Sciences
4301 Jones Bridge Rd
Bldg C, Rm C2027
Bethesda, MD 20814

**This PDF file includes:**

Figures S1 to S14

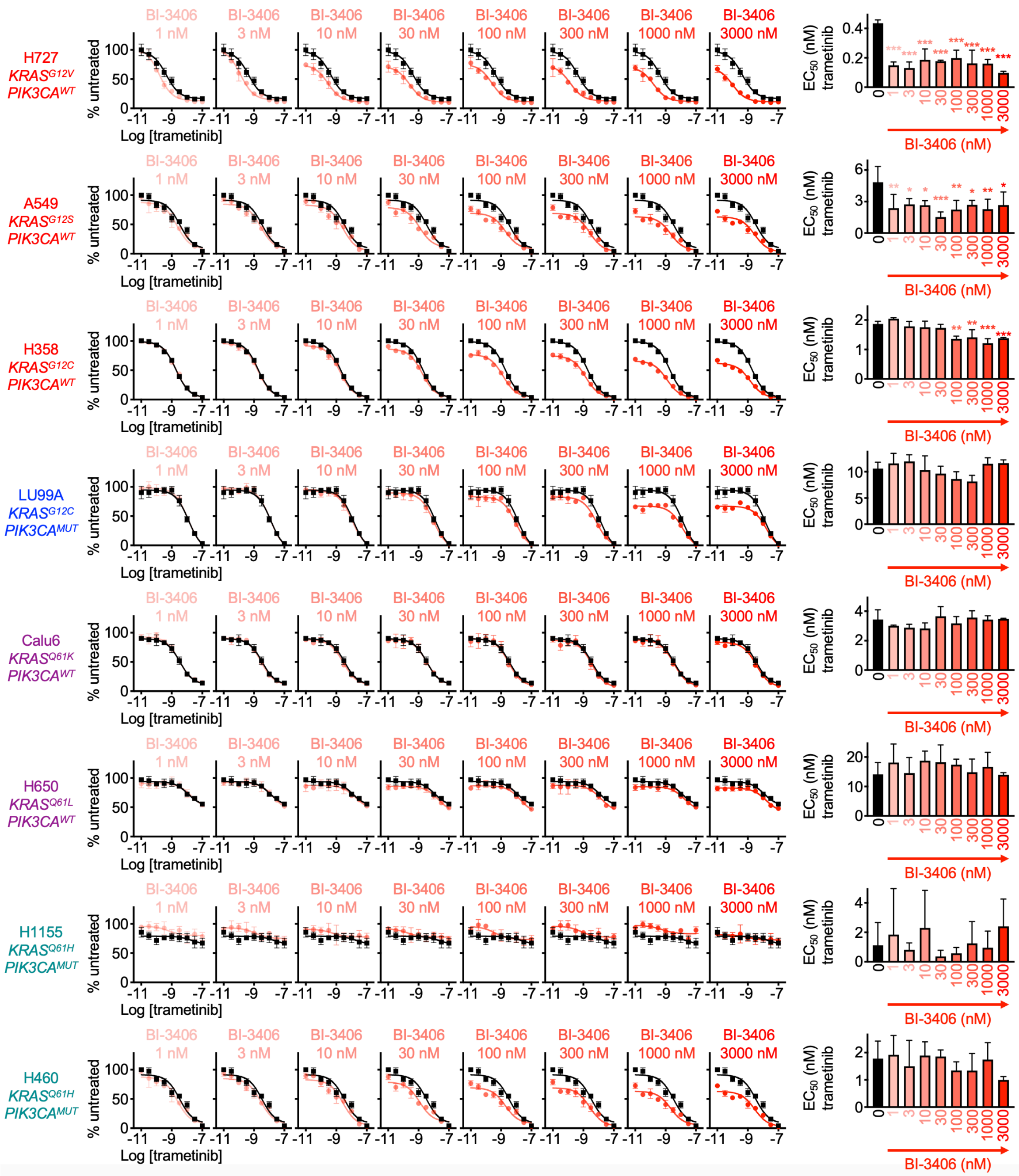


**Fig. S1** (related to Fig. 1 A). SOS1 inhibition sensitizes *KRAS*$^{G12}$/*PIK3CA*$^{WT}$ LUAD cells to the killing effect of trametinib. Trametinib dose response curves at each dose of BI-3406 (color) versus trametinib alone (black) indicating % cell viability (left) and $EC_{50}$ values (right) for *KRAS*-mutated LUAD cell lines treated with increasing (semilog) doses of trametinib ($10^{-10.5}$ – $10^{-7}$), BI-3406 ($10^{-9}$ – $10^{-5.5}$) or the combination of trametinib + BI-3406 under 3D spheroid culture conditions for 72 h. * $p<0.05$, ** $p<0.01$, *** $p<0.001$ vs no BI-3406 treatment.

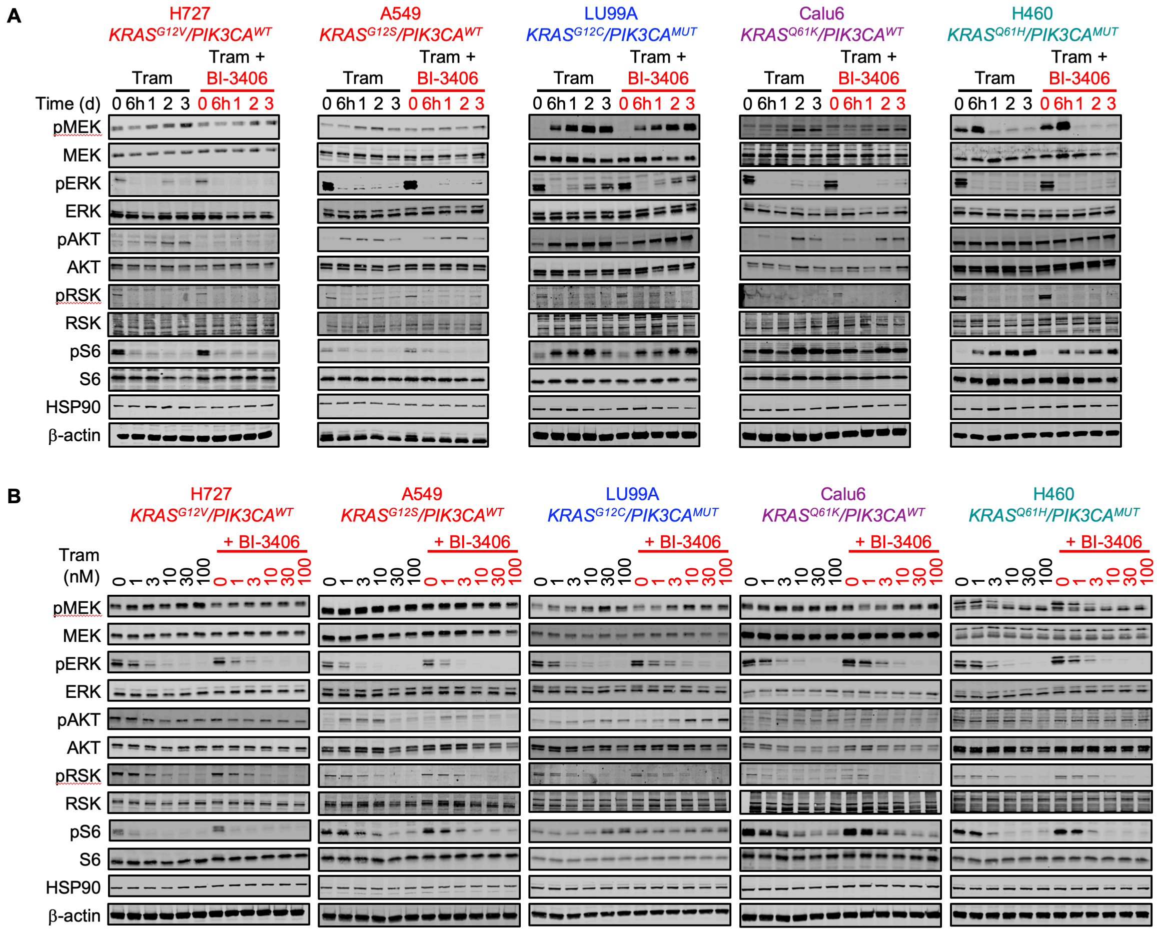


**Fig. S2** (related to Fig. 1 C). SOS1 inhibition increases the efficacy of and prevents rebound signaling to trametinib in *KRAS*$^{G12}$/*PIK3CA*$^{WT}$ LUAD cells. *(A)* Western blots of WCLs of 3D spheroid cultured *KRAS*$^{G12}$-mutated LUAD cell lines treated with trametinib (10 nM) ± BI-3406 (300 nM) for the indicated times. *(B)* Western blots of WCLs of 3D spheroid cultured *KRAS*$^{G12}$-mutated LUAD cell lines treated with increasing doses of trametinib ± BI-3406 (300 nM) for 24 h. Western blots are for pMEK, MEK, pERK, ERK, pAKT (Ser 473), AKT, pRSK, RSK, pS6, S6, HSP90, β-actin.

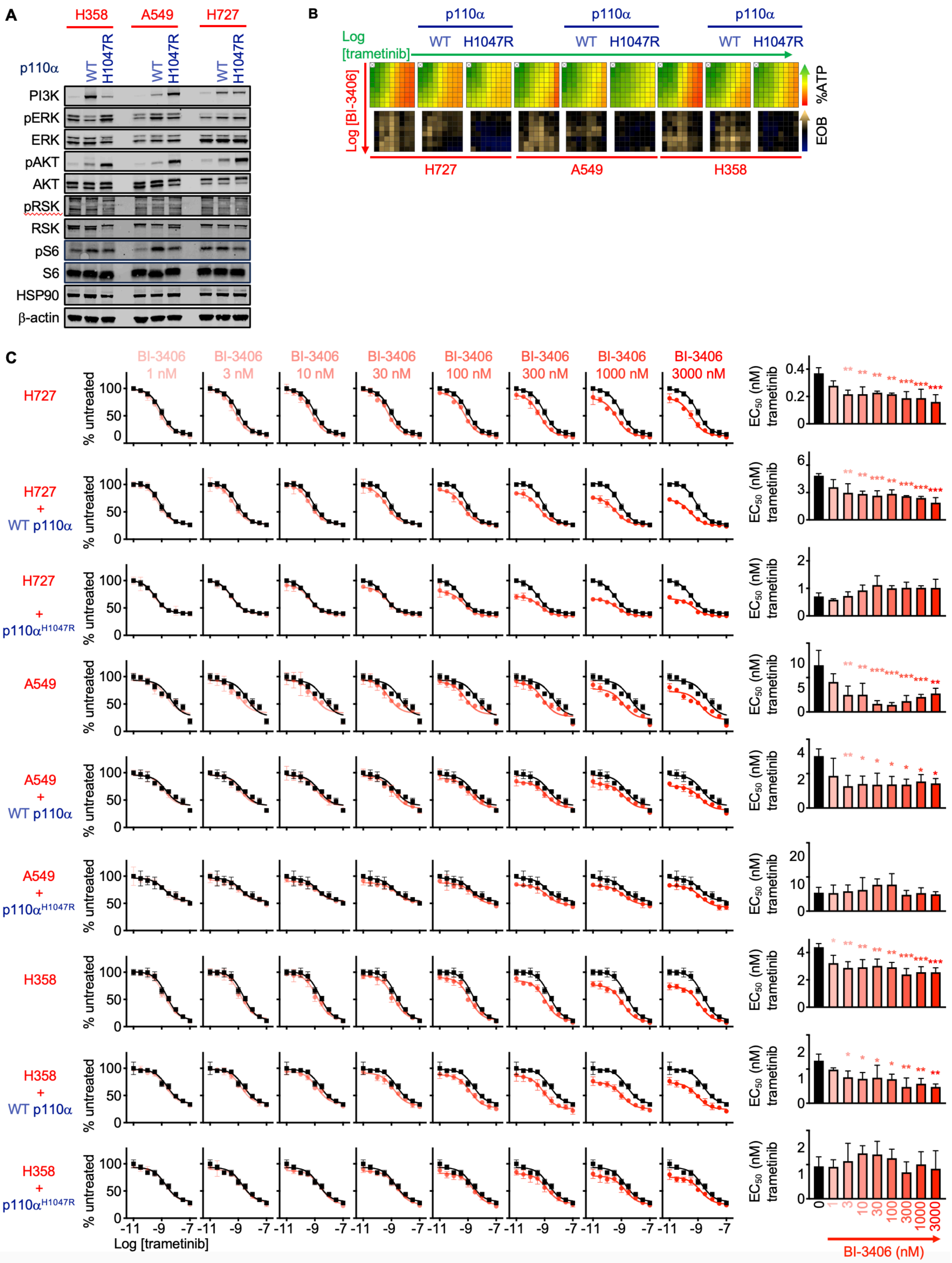


**Fig. S3** (related to Fig. 1 D). Expression of an activated p110α catalytic subunit inhibits SOS1 inhibitor-dependent sensitization of *KRAS*$^{G12}$/*PIK3CA*$^{WT}$ LUAD cells to the killing effect of trametinib. SOS1 inhibitor sensitivity. *(A)* Western blots of WCLs from the indicated LUAD cell lines expressing WT or H1047R mutant p110**α** catalytic subunit. Western blots are for pMEK, MEK, pERK, ERK, pAKT (Ser 473), AKT, pRSK, RSK, pS6, S6, HSP90, β-actin. *(B-C)* Heat map of cell viability *(B)* and trametinib dose response curves at each dose of BI-3406 (color) versus trametinib alone (black) indicating percent cell viability (left) and $EC_{50}$ values (right) *(C)* for *KRAS*$^{G12}$/*PIK3CA*$^{mut}$ LU99A cells treated with the indicated dose of the PI3K inhibitor copanlisib and increasing (semilog) doses of trametinib ($10^{-10.5}$ – $10^{-7}$), BI-3406 ($10^{-9}$ – $10^{-5.5}$) or the combination of trametinib + BI-3406 under 3D spheroid culture conditions for 72 h. * $p<0.05$, ** $p<0.01$, *** $p<0.001$ vs no BI-3406 treatment.

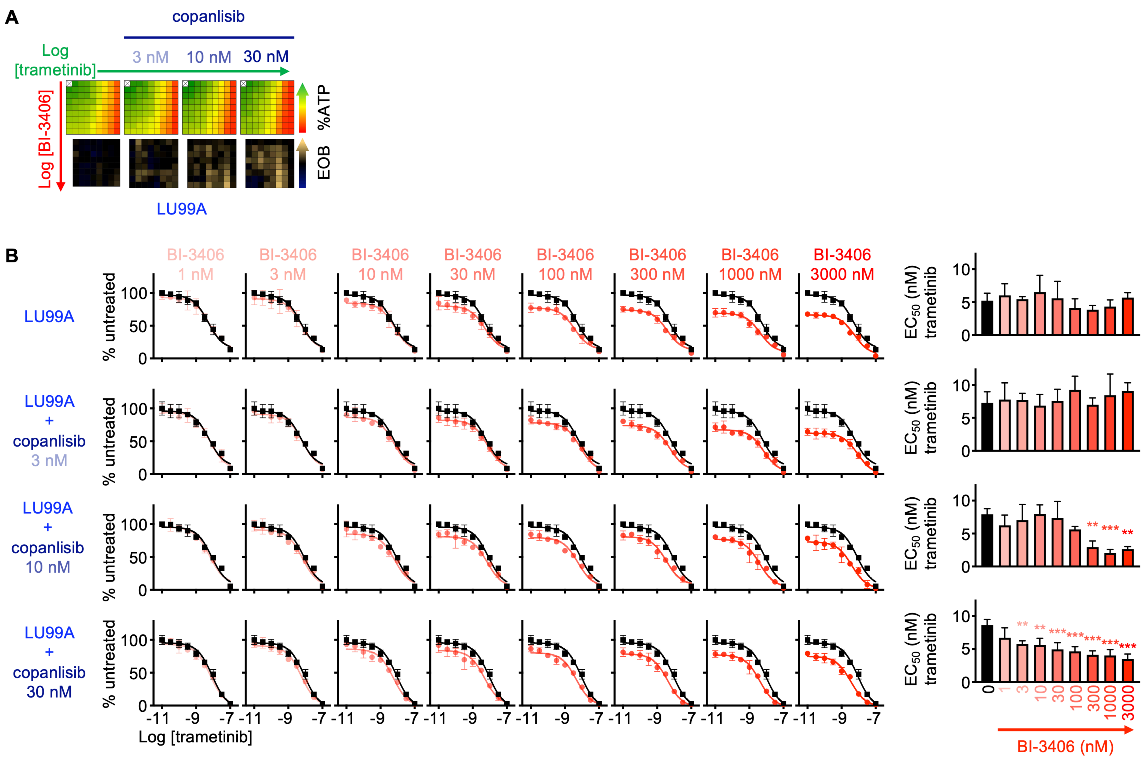


**Fig. S4** (related to Fig. 1 E). PI3K inhibition restores the ability of SOS1 inhibition to sensitize *KRAS*$^{G12}$/*PIK3CA*$^{MUT}$ LU99A cells to the killing effect of trametinib. *(A-B)* Heat map of cell viability *(A)* and trametinib dose response curves at each dose of BI-3406 (color) versus trametinib alone (black) indicating percent cell viability (left) and EC$_{50}$ values (right) *(B)* for *KRAS*$^{G12}$/*PIK3CA*$^{mut}$ LU99A cells treated with the indicated dose of the PI3K inhibitor copanlisib and increasing (semilog) doses of trametinib ($10^{-10.5}$ – $10^{-7}$), BI-3406 ($10^{-9}$ – $10^{-5.5}$) or the combination of trametinib + BI-3406 under 3D spheroid culture conditions for 72 hours. * $p<0.05$, ** $p<0.01$, *** $p<0.001$ vs no BI-3406 treatment.

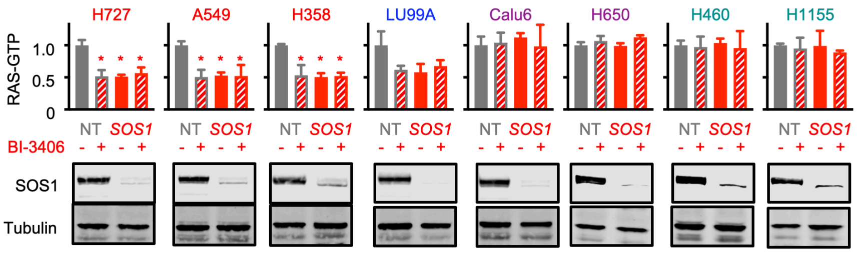


**Fig. S5** (related to Fig. 1 F). SOS1 ablation (via either BI-3406 treatment or *SOS1* KO) inhibits RAS activation in $KRAS^{G12}$-mutated cells. G-LISA was used to assess GTP-bound RAS in NT and *SOS1* KO $KRAS^{G12}$-mutated LUAD cells ± BI-3406 (300 nM). Western blots for SOS1 and β-actin showing *SOS1* KO are shown.

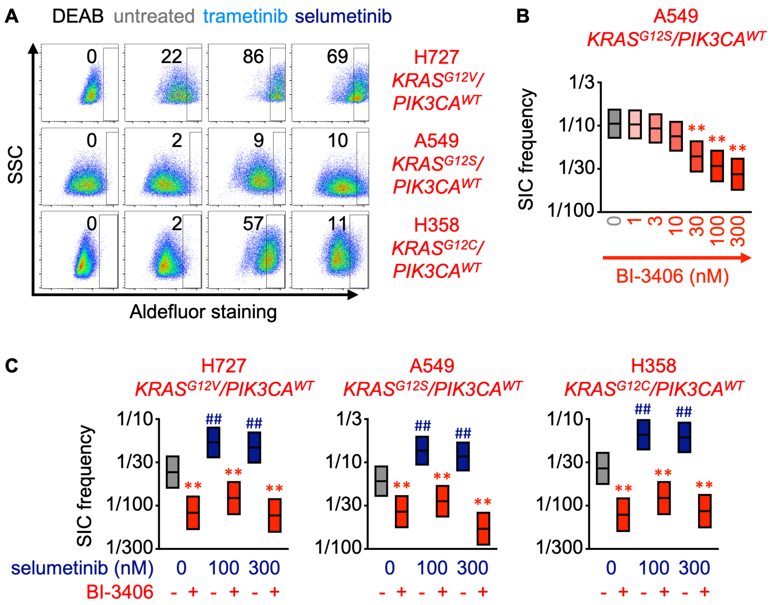


**Fig. S6** (related to Fig. 2 A through D). SOS1 inhibition prevents MEK inhibitor induced SIC outgrowth. *(A)* Aldefluor staining for ALDH enzyme activity in DEAB negative control (DEAB), untreated cells, or cells treated with 100 nM trametinib or selumetinib for 72 h for the indicated cell lines. H727 data are the same as in Fig. 2 A and are repeated here for comparison purposes. *(B)* SIC frequency from *in situ* ELDAs of A549 cells treated with the indicated doses of BI-3406. *(C)* SIC frequency from *in situ* ELDAs of the indicated cell lines pre-treated with selumetinib for 72 h to upregulate SICs, and then left untreated or treated with BI-3406. # p<0.05 vs untreated; ## p<0.01 vs. untreated for SIC upregulation by MEK inhibitor treatment vs. untreated controls. * p<0.05 vs untreated; ** p<0.01 for SIC inhibition by BI-3406 treatment compared to untreated controls. Data are representative from three independent experiments.

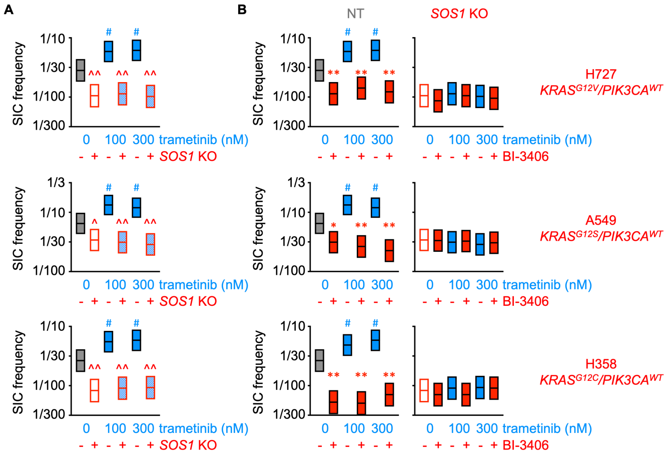


**Fig. S7** (related to Fig. 2 E). SOS1 KO inhibits trametinib-induced SIC upregulation in *KRAS*$^{G12}$-mutated *PIK3CA*$^{WT}$ LUAD cells. *(A)* SIC frequency from *in situ* ELDAs in the indicated NT and *SOS1* KO COAD cells pre-treated with trametinib for 72 h. *(B)* SIC frequency from *in situ* ELDAs in the indicated NT and *SOS1* KO COAD cells pre-treated with trametinib for 72 h to upregulate SICs, and then left untreated or treated with the SOS1 inhibitor BI-3406 to assess BI-3406 specificity. # p<0.05 vs untreated, vs. untreated for SIC upregulation by MEK inhibitor treatment vs. untreated controls. * p<0.05, ** p<0.01 for SIC inhibition by BI-3406 treatment compared to untreated controls. ^ p<0.05; ^^ p<0.01 for *SOS1* KO compared to untreated controls.

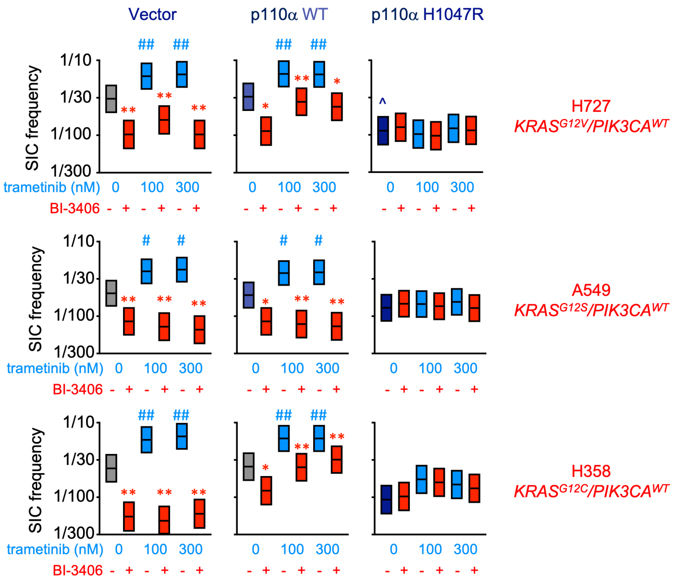


**Fig. S8** (related to Fig. 2 F). Constitutive PI3K/AKT pathway activation inhibits trametinib- and BI-3406-dependent SIC regulation. SIC frequency from *in situ* ELDAs in the indicated LUAD cell lines expressing WT or H1047R mutant p110α catalytic subunit vs vector control cells. # p <0.05 vs untreated, vs. untreated for SIC upregulation by MEK inhibitor treatment vs. untreated controls. * p<0.05, ** p<0.01 for SIC inhibition by BI-3406 treatment compared to untreated controls. ^ p<0.05; for p110α$^{H1047R}$ compared to untreated controls.

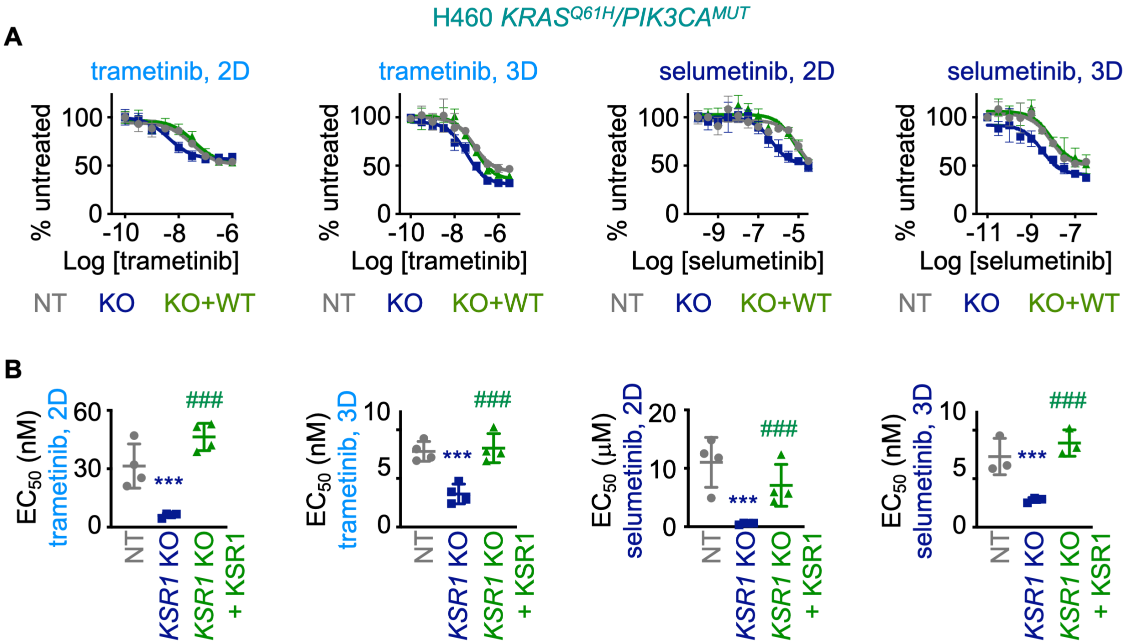


**Fig. S9** (related to Fig. 3 B and C). *KSR1* KO sensitizes *KRAS*$^{Q61H}$-mutated LUAD cells to the killing effects of MEK inhibition. *(A-B)* Single-dose response curves *(A)* and the $EC_{50}$ values *(B)* in NT, *KSR1* KO, and *KSR1* KO+*KSR1* cells H460 (*KRAS*$^{Q61H}$/*PIK3CA*$^{MUT}$) LUAD cells at 2D (anchorage-dependent) and 3D (anchorage-independent) conditions in cells treated with increasing doses of trametinib or selumetinib for 72 h. Data are averaged from three independent experiments. *** p<0.01 vs non-targeting controls; ### p<0.001 vs. *KSR1* KO.

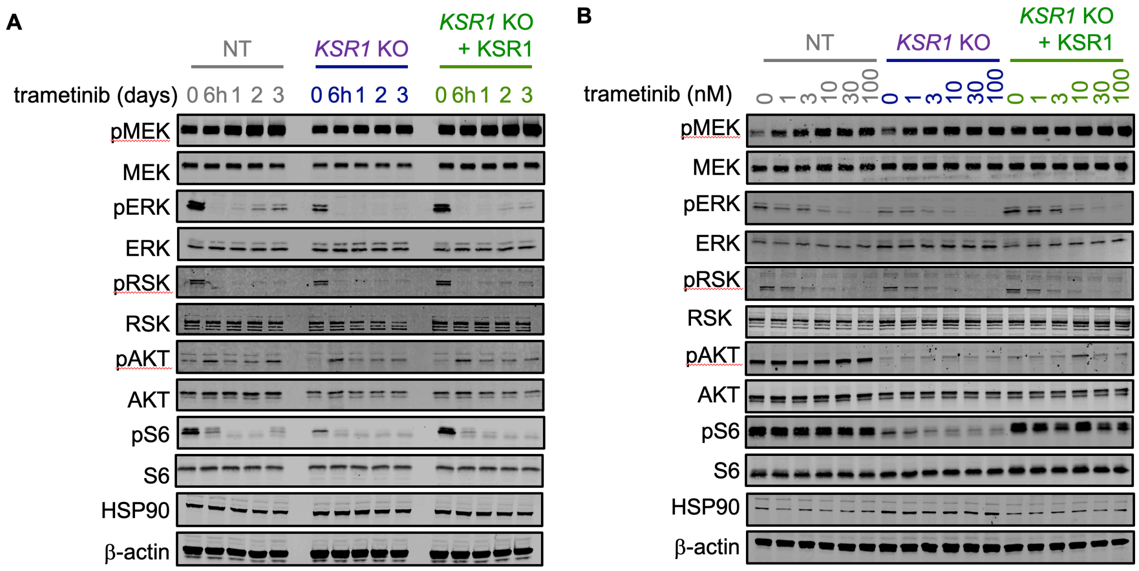


**Fig. S10** (related to Fig. 3 D and E). *KSR1* KO sensitizes H460 cells to the killing effects of trametinib. *(A-B)* Western blots of WCLs H460 *KSR1* KO an NT cells treated with trametinib (100 nM) for the indicated times *(A)* or with the indicated dose of trametinib for 24 h *(B)*. Western blots are for pMEK, MEK, pERK, ERK, pAKT (Ser 473), AKT, pRSK, RSK, pS6, S6, HSP90, β-actin.

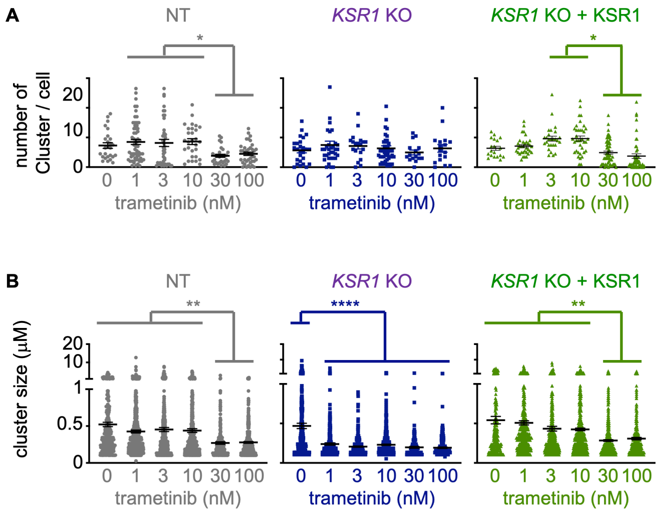


**Fig. S11** (related to Fig. 3 F and G). *KSR1* KO sensitizes H460 cells to the killing effects of trametinib. *(A-B)* Quantification of the number of MEK-ERK clusters from *(A)* and size of individual clusters (B) from proximity ligation assays in H460 cells treated with the indicated dose of trametinib for 24 h from Fig. 3 F The threshold for individual clusters was set to 0.1 μm$^2$. In (A) each individual point represents a cell, in (B) each individual point represents a MEK-ERK cluster. Data are quantified from >20 cells from three fields from three independent experiments.

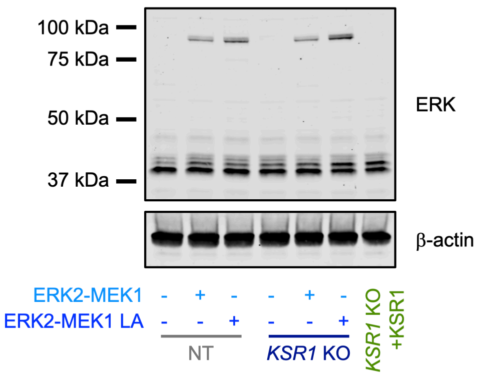


**Fig. S12** (related to Fig. 3 H). An ERK2-MEK1 fusion protein rescues clonogenicity in *KSR1* KO H460 cells Western blots of WCLs H460 *KSR1* KO an NT cells expressing a WT or nuclear localized (LA) YFP-tagged MEK2-ERK1 fusion protein. Western blots are for ERK and β-actin. The YFP-tagged fusion protein runs at ~90 kDa.

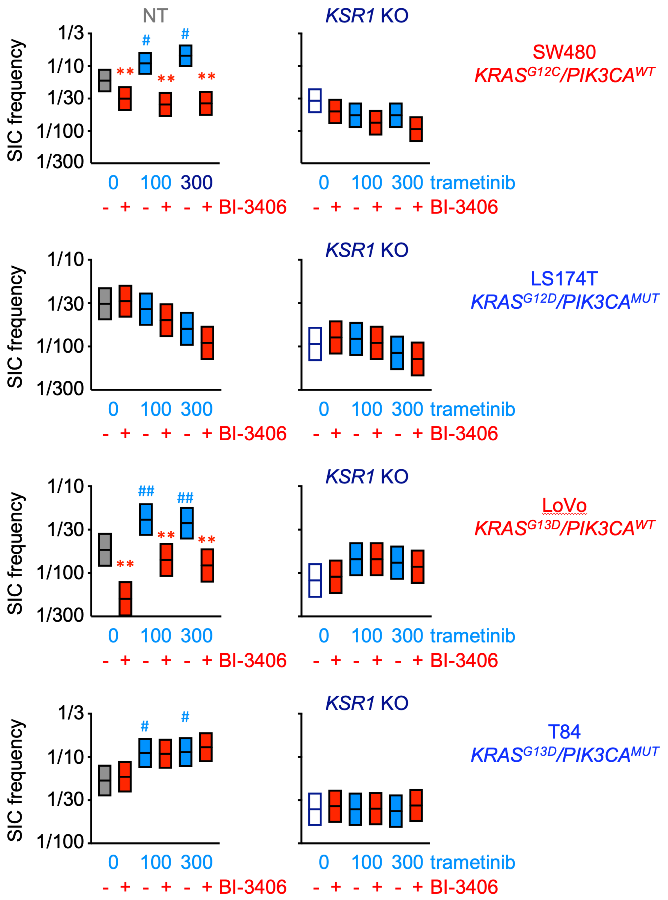


**Fig. S13** (related to Fig. 5). SOS1 inhibition does not enhance the effect of *KSR1* KO on basal and trametinib-induced SICs in *KRAS*-mutated COAD cells. SIC frequency from *in situ* ELDAs of the indicated NT and *KSR1* KO COAD cells pre-treated with trametinib for 72 h to upregulate SICs, and then left untreated or treated with the SOS1 inhibitor BI-3406. The *KRAS* and *PIK3CA* mutational status for each cell line is indicated. NT cells in the left column are from Fig. 4 and are shown here for comparison purposes. # $p<0.05$ vs untreated; ## $p<0.01$ vs. untreated for SIC upregulation by MEK inhibitor treatment vs. untreated controls. ** $p<0.01$ for SIC inhibition by BI-3406 treatment compared to untreated controls.

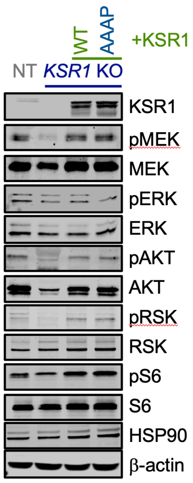


**Fig. S14** (related to Fig. 5 A). KSR1 regulation of TICs/SICs in COAD is dependent on interaction with ERK and relevant *in vivo*. Western blot for KSR1, pMEK, MEK, pERK, ERK, pAKT (Ser 473), AKT, pRSK, RSK, pS6, S6, HSP90, β-actin in WCLs of HCT116 (*KRAS*$^{G13D}$/*PIK3CA*$^{mut}$ NT, *KSR1* KO, *KSR1* KO + *KSR1* addback, and *KSR1* KO+ERK-binding mutant KSR1 (KSR1$^{AAAP}$) addback cells.
